# Single-cell analysis of Crohn’s disease lesions identifies a pathogenic cellular module associated with resistance to anti-TNF therapy

**DOI:** 10.1101/503102

**Authors:** JC Martin, G Boschetti, C Chang, R Ungaro, M Giri, LS Chuang, S Nayar, A Greenstein, M. Dubinsky, L Walker, A Leader, JS Fine, CE Whitehurst, L Mbow, S Kugathasan, L.A. Denson, J. Hyams, JR Friedman, P Desai, HM Ko, I Laface, Guray Akturk, EE Schadt, S Gnjatic, A Rahman, M Merad, JH Cho, E Kenigsberg

## Abstract

Clinical benefits to cytokine blockade in ileal Crohn’s disease (iCD) have been limited to a subset of patients. Whether cellular and molecular heterogeneity contributes to variability in treatment responses has been unclear. Using single cell technologies combining scRNAseq, CyTOF and multiplex tissue imaging, we mapped the cellular landscape of inflamed ileum lesions, adjacent non-inflamed ileum and matched circulating blood cells of iCD patients. In inflamed tissues, we identified a pathogenic module characterized by an inflammatory mononuclear phagocyte (Inf.MNP)-associated cellular response organized around inflammatory macrophages and mature dendritic cells in a subset of iCD patients. We confirmed the Inf.MNP-associated cellular response in 4 independent iCD cohorts (n=441) and showed that presence of this pathogenic module at diagnosis correlated with primary resistance to anti-TNF therapy. Single cell mapping of iCD tissues identifies a complex cellular signature of anti-TNF resistance thereby revealing novel biomarkers of treatment response and tailored therapeutic opportunities.

## Introduction

Inflammatory bowel disease (IBD), which comprises Crohn’s disease (CD) and ulcerative colitis, is characterized by intermittent chronic inflammation of the gastrointestinal tract leading to bowel damage and disabilities (Torres et al., 2017). IBD results from a complex interplay between westernized lifestyle-associated environmental factors and genetic susceptibilities, culminating in uncontrolled immune responses against luminal triggers (Kaser et al., 2010). Genome-wide association studies (GWAS) have identified more than 200 IBD-associated loci coherently organized into regulatory networks enriched for immune and inflammatory processes (Jostins et al., 2012; Liu et al., 2015). In order to design new drugs targeting immune mediators specifically involved in IBD lesions, numerous efforts combining human tissue analyses and rodent colitis models, have attempted to dissect the key cellular and molecular modules of intestinal inflammation (Neurath, 2017; de Souza and Fiocchi, 2015). The observation that therapeutic responses to immune biotherapies have constantly been limited to a subset of patients, however, suggests that similar clinical phenotypes can emerge from distinct inflammatory signatures (Abraham et al., 2016; Danese et al., 2016). Current approaches restricted to well-established antibody panels based on prior knowledge preclude the identification of novel pathogenic cell populations that span the diseased intestine. Recent significant advances of single-cell sequencing technologies now allows the characterization of human lesional tissues at high-resolution (Jaitin et al., 2014; Macosko et al., 2015; Klein et al., 2015; Zheng et al., 2017a; Azizi et al., 2018). In this study, we sought to map the cellular landscape of inflamed ileum lesions, adjacent non-inflamed ileum and matched circulating blood cells of iCD patients to help dissect disease heterogeneity between patients and identify the underlying cellular and molecular events that may control distinct disease outcome and response to treatment.

## Results

### High-resolution cell-type mapping of inflamed and uninflamed ileums in Crohn’s disease

Lamina propria cells were isolated from paired uninflamed and inflamed biopsies obtained from surgically resected ileums of 11 iCD patients (**Figure 1A; Table S1**). Single-cell transcriptomes were isolated from 22 ileal specimen, and Unique Molecular Identifiers (UMIs) counts matrices were generated (Zheng et al., 2017b) (**Table S2; supplemental methods**). After exclusion of epithelial and red blood cells as well as cells not passing quality controls (**Figures S1A-C**), 82,417 lamina propria cells from the 22 samples (**Figure S1D**) were clustered jointly. Based on our previous work, we used an expectation maximization (EM)-like clustering algorithm, which iteratively learns the gene expression profiles of the different cell populations while estimating batch-specific background noise rates (**Figures S1E-I; supplemental methods**) (Jaitin et al., 2014; Paul et al., 2015). Our clustering analysis revealed 47 clusters with variable number of cells (155-6,940 cells) (**Figure S1J**) and UMIs counts per cell (**Figure S1K)**. All clusters included cells from multiple patients, suggesting that cells were grouped according to shared lamina propria-induced program rather than patient specificity (**Table S3**).

**Figure 1.**
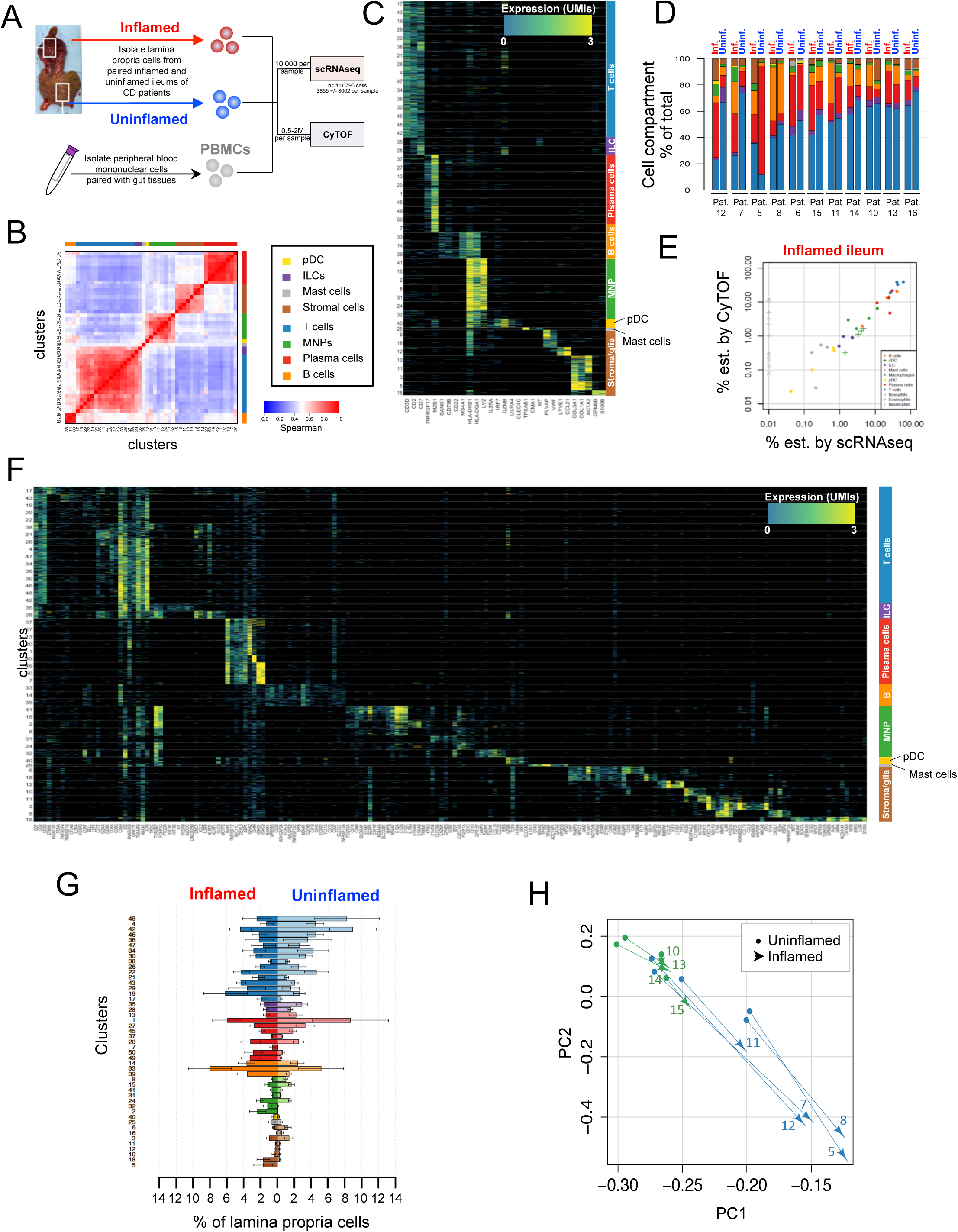
High-resolution cell-type mapping of inflamed and uninflamed ileums in Crohn’s disease. **(A)** Workflow showing the processing of freshly collected blood and surgical resections, including paired inflamed and uninflamed tissues of ileal Crohn’s disease patients for scRNAseq and CyTOF. (**B**) Grouping scRNAseq clusters by similarity. Heatmap showing Pearson correlation coefficients between the log average expression profiles of pairs of clusters. Clusters were reordered by hierarchical clustering. (**C)** Expression of key cell-type markers across the 8 cell compartments. Stacked rows correspond to single-cells down-sampled to 2000 UMIs/cell and columns to selected marker genes for which UMI counts are color-coded. 100 randomly selected cells per cluster cell are visualized. Clusters are demarcated by gray bars and grouped by lineage according to marker genes. **(D)** Percentages of cellular lineages in individual samples included in the scRNAseq analysis. **(E)** CyTOF validation of frequencies estimated by scRNAseq. Frequencies of the 7 immune lineages as estimated by CyTOF vs. scRNAseq in 4 inflamed iCD ileums. **(F)** High-resolution characterization of cellular subtypes identity. Same map as shown in **C** extended with genes showing high variability between clusters and demonstrating the diverse cell states captured by scRNAseq. **(G)** Percentages of individual scRNAseq clusters in 9 inflamed and 9 uninflamed ileums (mean ± SEM). **(H)** Principle component analysis of cell subtype composition. First two components provided by principal component analysis of adjusted cell subtype frequencies (frequencies were normalized by cellular lineages). Dots and arrowheads correspond to uninflamed and inflamed ileums respectively of the different patients indicated by their IDs. Paired uninflamed and inflamed ileal tissues are connected by a line.

Hierarchical clustering of averaged expression profiles grouped the 47 clusters into 8 major cellular compartments (**Figure 1B**), separating stromal/glial cells from 7 distinct immune cell lineages consisting of T cells, innate lymphoid cells (ILCs), B cells, plasma cells, mononuclear phagocytes (MNP), plasmacytoid dendritic cells (pDCs) and mast cells (**Figure 1C**). Frequency estimation of the major immune cellular compartments identified by scRNAseq was validated by pairing single-cell analysis with cytometry by time-of-flight (CyTOF) (**Figures 1D&E, S2A**). Importantly, scRNAseq sensitivity to estimate rare subpopulations with frequencies as low as 0.2% in individual samples such as pDCs, was verified (**Figures 1E and S2A&B**). In agreement with previous studies, granulocyte populations were not captured by 10x Chromium scRNAseq, while they were identified by CyTOF (**Figure S2C**) (Ordovas-Montanes et al., 2018). Within the 8 major cellular compartments, deeper transcriptomic characterization of the 47 clusters revealed distinct cell subtypes and/or cell states (**Figure 1F**), many of which were significantly different between inflamed and uninflamed ileum (**Figure 1G**). Principal component analysis (PCA) of normalized subtype frequencies demonstrated that 84% of their variance could be explained by only two principal components, suggesting that while inflamed tissues followed similar trend of changes from their paired uninflamed counterpart, a second layer of variability existed between inflamed ileums which separated the patients into two subgroups (**Figure 1H**).

### Two distinct inflammatory MNP populations reside in ileal Crohn’s disease lesions

Risk allele gene expression association established the MNP compartment as a central component of CD pathophysiology (Peters et al., 2017). Accordingly, MNP clusters were highly variable between inflamed and uninflamed ileum (**Figure 1G**). MNP clusters included dendritic cells (DC) expressing the hallmark receptor *FLT3*, and macrophages expressing the “core macrophage gene signature” that includes *MERTK*, *CTSC*, *CTSD*, *GLUL* and *PLD3* (Gautier et al., 2012), in addition to known human macrophage markers i.e. *CD14*, *CD68* and *FCGR3A* (CD16a) (**Figure 2A**). Analyses of gene modules with strong covariance of expression within the myeloid cells (**Figure S3A, Table S4**) further revealed two subtypes of macrophages exhibiting tissue-resident and infiltrating macrophage signatures respectively. The gut tissue-resident macrophage subset expressed high levels of the macrophage transcription factors (TF) *MAFB* and M-CSF receptor (*CSF1R*) (Lavin et al., 2014), genes required for the clearance of cellular debris (*DNASE1L3* and complement 1q genes *C1QA*, *C1QB*, *C1QC*) (Sisirak et al., 2016), immune modulation (e.g. the B7 family related-member *VSIG4*), and resolution of inflammation (*MRC1*) (Lee et al., 2002; Li et al., 2017) (**Figure 2B**). Further analysis showed the restricted expression of several genes like *FOLR2*, *FUCA1*, *GPNMB* and *LILRB5* to one of the two clusters of gut tissue-resident macrophage, possibly indicative of different metabolic states and/or niche within the ileum. The infiltrating macrophage subset likely corresponded to inflammatory monocyte-derived macrophages (infl. macs), as supported by the expression of the TF NF-κB. Accordingly infl. macs expressed a plethora of inflammatory mediators including the two major CD therapeutic targets TNF and IL-23, and the IBD biomarker calprotectin (*S100A8* and *S100A9*) (**Figure 2B**) (Konikoff and Denson, 2006). Within the DC compartment, we identified transcriptomic signatures corresponding to the well-characterized CD1c^+^/DC2 and CD141^+^/DC1 (Guilliams et al., 2016; See et al., 2017; Villani et al., 2017). A third DC cluster expressing the *CD207* (langerin) together with *CD209* (DC-SIGN) and other mononuclear phagocytic genes including C1Qs, *SEPP1* and *CD163*, likely corresponded to monocyte-derived DC. The last DC cluster was enriched in transcripts usually present in migratory DC on their way to the tissue draining lymph node (DLN), which included the activation marker DC-LAMP (*LAMP3*) together with co-stimulatory molecules (*CD40*, *CD86* and *CD274* aka. PD-L1) and the chemokine receptor CCR7 (**Figure 2C**). The myeloid cell diversity observed in the CD ileum was validated by CyTOF analysis (**Figure S3B-F**), and both infl. macs and mature DCs were identified as strongly enriched in inflamed vs. uninflamed tissues by scRNAseq (11-fold and 7-fold respectively) and CyTOF (**Figures 2D-E**).

**Figure 2.**
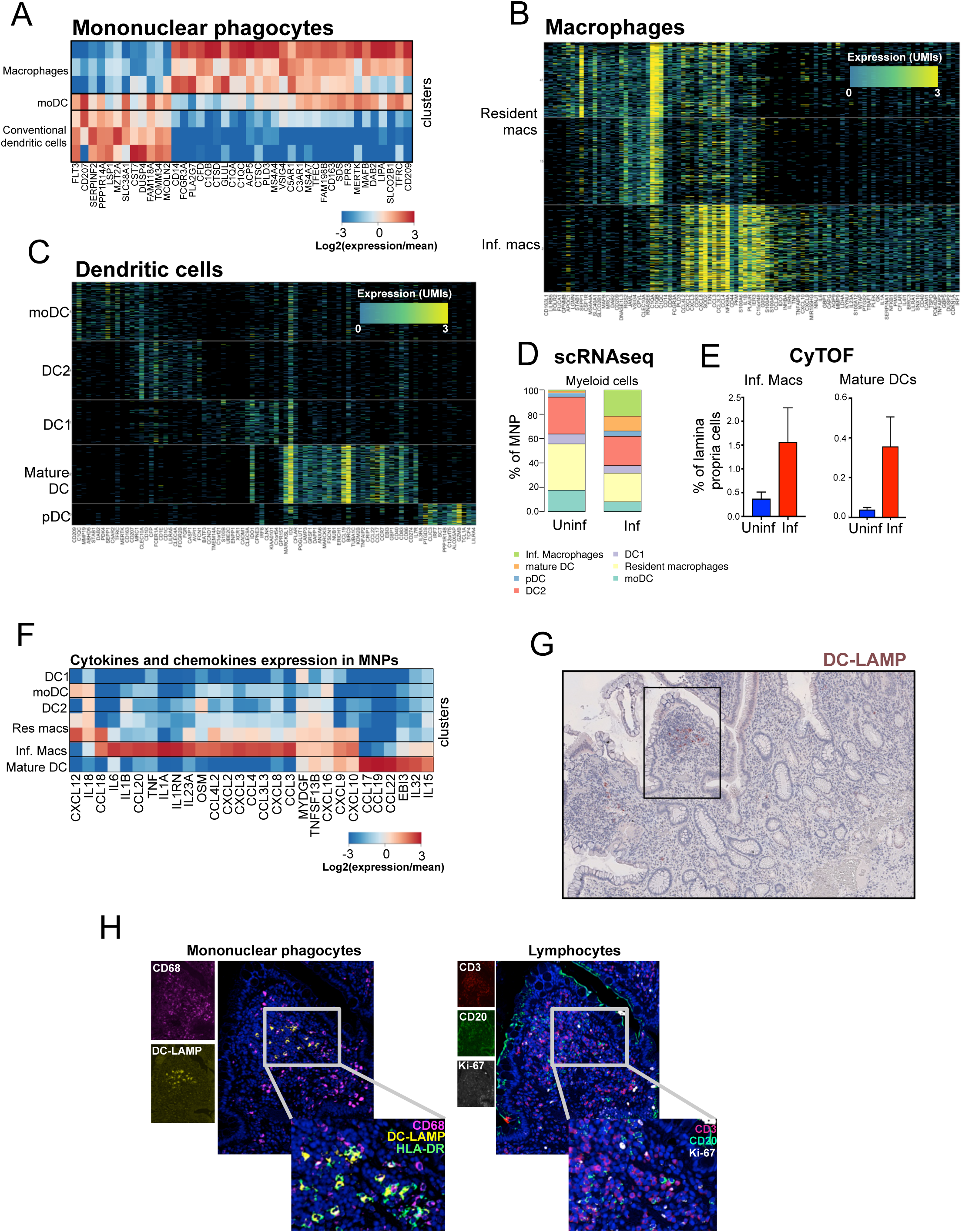
Two distinct inflammatory MNP populations in ileal Crohn’s disease lesions. **(A)** Distinct expression profiles identify macrophages and dendritic cells. Heat-map showing relative expression values of genes (columns) enriched in macrophages or dendritic cells clusters (rows). **(B-C)** Distinct transcriptional programs between MNP subtypes. Heat-maps showing color-coded down-sampled UMI counts of variable genes between macrophage clusters (B) and between dendritic cell clusters (C). Clusters are demarcated by gray bars. For each cluster, 300 cells were randomly selected and down-sampled to 2000 UMIs/cell. **(D)** MNP subtypes composition in inflamed and uninflamed regions. Shown are stacked frequencies of each MNP subtypes divided by the total MNP frequency estimated by scRNAseq in uninflamed or inflamed ileum. **(E)** CyTOF validation of inflammatory macrophages and mature DC expansion in inflamed ileum. Bar graphs showing average frequencies of indicated populations as determined by CyTOF analysis of uninflamed and inflamed ileums (n=7 CD patients; mean ± SEM). **(F)** Heat-map showing relative expression of cytokines and chemokines detected at significant levels in MNP clusters. **(G)** Representative staining of DC-LAMP^+^ mature DC in the inflamed lamina propria. The black square identifies the region with a mature DC-centered lymphoid aggregate presented at higher magnification in (H). (**H**) Representative staining of a HLA-DR^+^ DC-LAMP^+^ mature DC-centered lymphoid aggregate in inflamed lamina propria.

Distinct patterns of cytokines and chemokines expression between infl. macs and mature DC suggested the former as major organizers of pathogenic innate immune response in ileal lesions (**Figure 2F**). Infl. macs preferentially expressed mediators of acute inflammatory responses, including neutrophil-attracting chemokines (CXCL2, CXCL3, CXCL8), as well as cytokines essential for ILCs and stromal cell activation like IL-23, IL-6, IL1β, IL-1α and TNF (Neurath, 2014; Powell et al., 2015). Accordingly, infl. macs also expressed the highest levels of oncostatin M (OSM), recently identified as an additional major contributor to pathogenic stromal activation in IBD lesions (West et al., 2017). In contrast, mature DC were enriched in adaptive lymphocyte-attracting chemokines including CCL17, CCL22 and CCL19, suggesting their potential role in the recruitment, activation, expansion and spatial organization of adaptive immune responses in iCD lesions (**Figure 2F**). The presence of a large cluster of mature DC in the inflamed ileum was unexpected as maturation is known to promote DC migration from the tissue to its DLN (Banchereau and Steinman, 1998). Using multiplexed immunohistochemical consecutive staining on a single slide (MICSSS) that we recently developed (Remark et al., 2016), we confirmed that infl. macs and mature DC both localized in the inflamed lamina propria **(Figures 2G and S3G**). In contrast to infl. macs, however, mature DC almost always colocalized with T and B lymphocytes forming dense lymphoid aggregates (**Figure 2H**). The high rate of Ki-67^+^ cycling lymphocytes further supported the local activation of adaptive T and B lymphocyte responses within the mature DC-centered lymphoid aggregates (**Figure 2G**). Importantly, while morphologically similar, mature DC-centered lymphoid aggregates differed from other T cell-dense areas surrounded by resident macrophages but devoid of B cells and mature DC (**Figure S3H**). Thus, we identified infl. macs and mature DC as two major populations of inflammatory MNP with distinct transcriptomic programs and spatial distribution, suggesting specialized contribution to pathogenic inflammation in iCD lesions.

### Identification of a cellular-associated module centered around inflammatory MNP in ileal Crohn’s disease lesions

Interestingly, correlation analysis of cell subtype frequencies defined by the scRNAseq clustering revealed that mature DC and infl. macs co-segregated together with specific lymphocyte and stromal cell populations in inflamed tissues (**Figure 3A**). Consistent with possible local induction of adaptive responses suggested by the mature DC-centered lymphoid aggregates observed in the inflamed lamina propria, lymphocyte populations preferentially associated with infl. macs and mature DC included highly activated cycling T cells, effector memory cytotoxic T cells (CTLs) and central memory T cells (T_CM_) with a Th1/17 cytokine/chemokine expression profile (**Figures 3B, S4A-E, and supplemental methods**) as well as plasmablasts and plasma cells (PC), which in contrast to IgA-producing tissue-resident PC, produced inflammatory IgG immunoglobulins (**Figures S4F&G and supplemental methods**). Group 1 ILCs, including NK cells and ILC1, (**Figure S4H and supplemental methods**) and regulatory T cells (Tregs) also co-segregated with the inflammatory MNP populations. Lymphocytes associated with infl. macs and mature DC were largely enriched in inflamed vs. uninflamed ileum, consistent with the pathogenic nature of the inflammatory MNP-associated immune signature (**Figure 3C)**. Finally, infl. macs and mature DC were also associated with two distinct populations of activated stromal cells, referred as ACKR1^+^ (aka. DARC) endothelial cells and activated fibroblasts (**Figures 3A, 3D&E and supplemental methods**).

**Figure 3.**
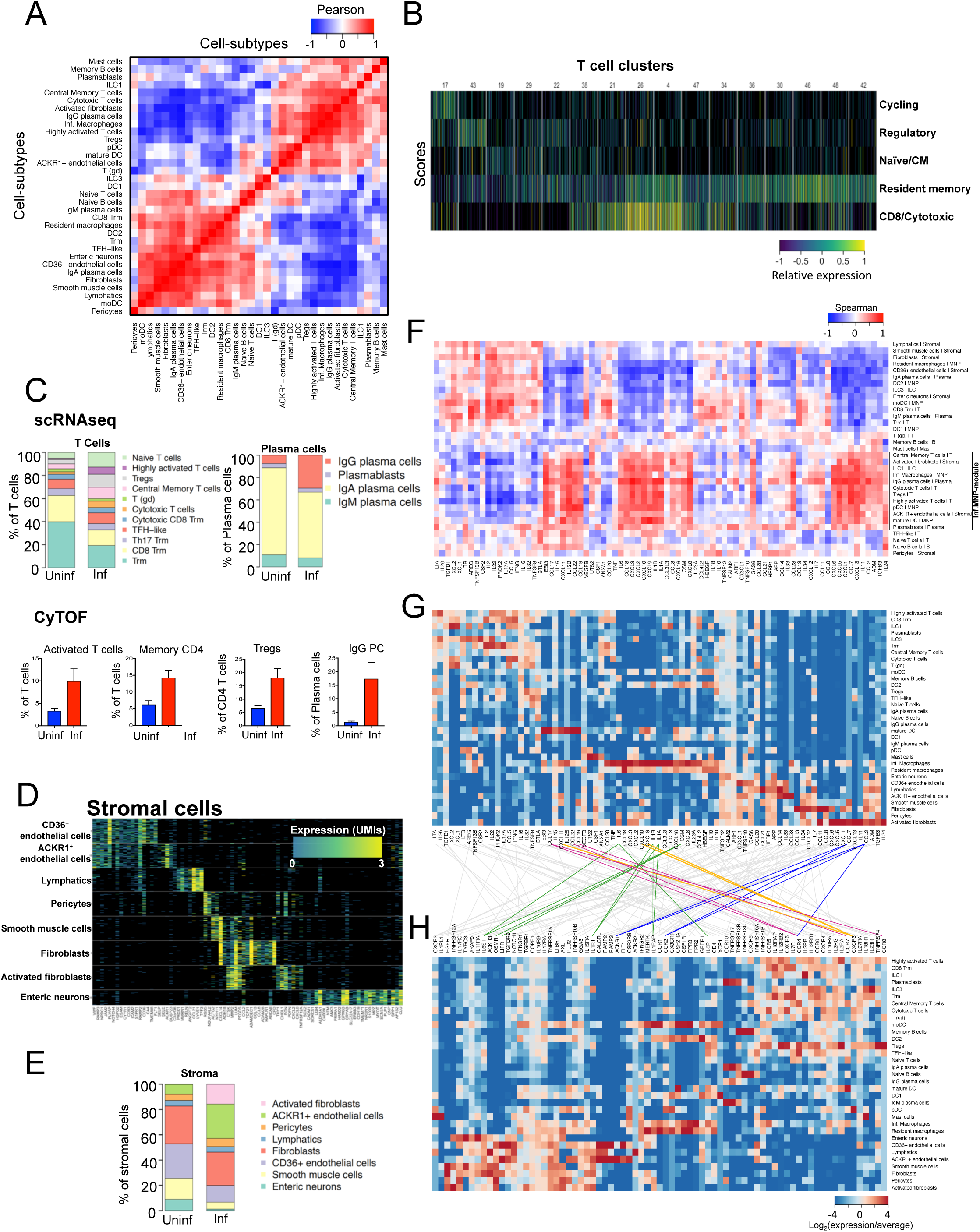
Identification of a cellular-associated module centered around inflammatory MNP in ileal Crohn’s disease lesions. **(A)** Co-segregation of cell-subtypes across inflamed tissues. Pearson correlation between the lamina propria cell subtypes in CD inflamed ileums based on their adjusted frequencies (n=9). Subtypes are reordered by hierarchical clustering. **(B)** Heat-map showing the relative expression of 5 transcriptional scores (rows) in single T cells (columns). Scores are integrating expression of multiple highly correlated genes (supplemental methods). Cell clusters are demarcated by gray bars. **(C)** Different composition of T cell and plasma cell (PC) subtypes in inflamed vs. uninflamed ileum. Top: Stacked frequencies of each T cell or PC subtypes divided by the respective total compartment frequency estimated by scRNAseq in uninflamed or inflamed ileum. Bottom: bar graphs of average frequencies of indicated populations as determined by CyTOF analysis of uninflamed and inflamed ileums (n=7 CD patients; mean ± SEM). **(D)** Stromal and glial diversity in inflamed and uninflamed ileum. Heatmap showing color-coded down-sampled UMI counts of variable genes (columns) between single-cells (rows) of different stromal subtypes. For each cluster, 100 cells were randomly selected and down-sampled to 2000 UMIs/cell. **(E)** Differences between stromal/glial composition in inflamed vs. uninflamed ileum. Frequencies of individual subtypes divided by the total stromal/glial cells frequency in uninflamed or inflamed ileum. **(F)** Heatmap showing the correlation between the frequencies of each cellular subtype in CD tissues (rows) and the total cytokine/chemokine expression over all other subtypes (columns). Subtype frequency is divided by the total frequency of its cellular compartment. **(G-H)** Ligand-receptor expression map. Heat-maps showing the average expression per cell subtype (rows) of indicated cytokines/chemokines (G) and their receptors (H). Established ligand-receptor pairs are connected with lines. Colored lines highlight pairs relevant to the Inf.MNP-module. Pink: mature DC-T_CM_; Yellow: Inf.MNP-activated T cells/CTLs/pDC; Green: infl. macs**-**stromal cells; Blue: activated fibroblasts-ACKR1^+^ endothelial cell.

To assess whether communications in the inflammatory MNP pathogenic signature could preferentially involve specific secreted mediators, we analyzed the correlations between the abundance of each cellular subtype and the total expression of individual cytokines and chemokines across all other cellular subtypes. Remarkably, we identified a specific set of cytokines/chemokines uniquely connected with the inflammatory MNP-associated response (**Figure 3F**). Mature DC-derived CCL17, CCL22 and CCL19 were correlated with these subtypes and combined expression of their respective receptors CCR4 and CCR7 was the highest for TCM, reflecting potential privileged and pathogenic interactions between mature DC and T_CM_ in mature DC-centered lymphoid aggregates. Concordantly, the cytokines/chemokines set also comprised CXCL9, CXCL10 and CXCL11, expressed by infl. macs and/or mature DC, which all bind to CXCR3, highly expressed by activated T cells, CTLs and Tregs, as well as by pDC (**Figures 3G&H**). Infl. macs-derived OSM, IL-1β and IL-1α were also identified and stromal cells expressed the highest levels of their receptors. Accordingly, endothelial cells and fibroblasts associated with the inflammatory MNP were activated (**Figure 3A**), and the latter expressed several cytokines and chemokines, including CXCL1, CXCL5, CXCL6, CCL2 and IL-11, inducible upon OSM/IL-1β exposure of colonic stromal cells in vitro and all identified in the Inf.MNP-associated cytokine/chemokine set (Bamba et al., 2003; Okuno et al., 2002; West et al., 2017) (**Figure 3F**). Altogether, these data unraveled the existence of a pathogenic response in the inflamed lamina propria, organized around mature DC-centered lymphoid aggregates and infl. macs, which was driven by a unique cytokine/chemokine program. The inflammatory MNP-associated response involved specific adaptive immune populations including activated T cells, Th1/17 T_CM_, effector memory CTLs and IgG-plasma cells, as well as activated stromal populations of fibroblasts and ACKR1^+^ endothelial cells. We coined the name inflammatory MNP-associated cellular module (Inf.MNP-module) to refer to it thereafter.

### The Inf. MNP-module is enriched in a subset of ileal Crohn’s disease patients

Quite strikingly, we found that the Inf.MNP-module was prominently enriched in the inflamed lamina propria in one of the two subgroups of patients identified by the PCA analysis in our cohort (**Figures 1H&4A**), and validated these results at the protein levels using CyTOF analysis (**Figure 4B**). Importantly, pathology severity, disease duration before surgery, and blood markers of inflammation at time of surgery were similar between patients with or without the Inf.MNP-module (**Figure S5A**). Interestingly, activated fibroblasts in the Inf.MNP-module expressed the chemokines CCL2 and CCL7 which mediate CCR2^+^ inflammatory blood classical monocytes transcytosis in inflamed tissues through the binding to ACKR1 on endothelial cells (Pruenster et al., 2009). scRNAseq clustering analysis of 42,983 peripheral blood mononuclear cells (PBMCs) matched to intestinal samples (**Figure S5B**) revealed that increased infl. macs frequencies in disease lesions was associated with a depletion of circulating CCR2^+^ classical monocytes in the blood of patients enriched for the Inf.MNP-module (**Figure 4C**), suggesting that blood CCR2^+^ monocytes likely contributed to the infl. macs pool through CCL2/CCL7 ACKR1^+^-mediated recruitment in the inflamed ileum. Inf.MNP-module high patients, however had significantly lower IL-2 expression in inflamed tissues (**Figure 4D**). We identified Trm as the main source of IL-2 in ileal tissues (**Figure 3G**), and we confirmed lower Trm-derived IL-2 in Inf.MNP-module high patients (**Figure 4D**). Since IL-2 is a major cytokine maintaining FOXP3 expression and immune suppressive function in Tregs (Rubtsov et al., 2010), these results suggested that Tregs were likely defective in lesions enriched in the Inf.MNP-module. Accordingly, while Tregs expanded in the inflamed mucosa of both subgroups, differential expression analysis revealed lower expression of *FOXP3* (Log2FC= −0.48, adj. p-value = 0.016) and FOXP3-targets (FDR-adjusted p value<10^−10^) (**Figure 4E, Table S5**) (Zheng et al., 2007) in Tregs from Inf.MNP-module high patients, suggesting that defective Tregs could contribute to the pathogenicity of the Inf.MNP-module.

**Figure 4.**
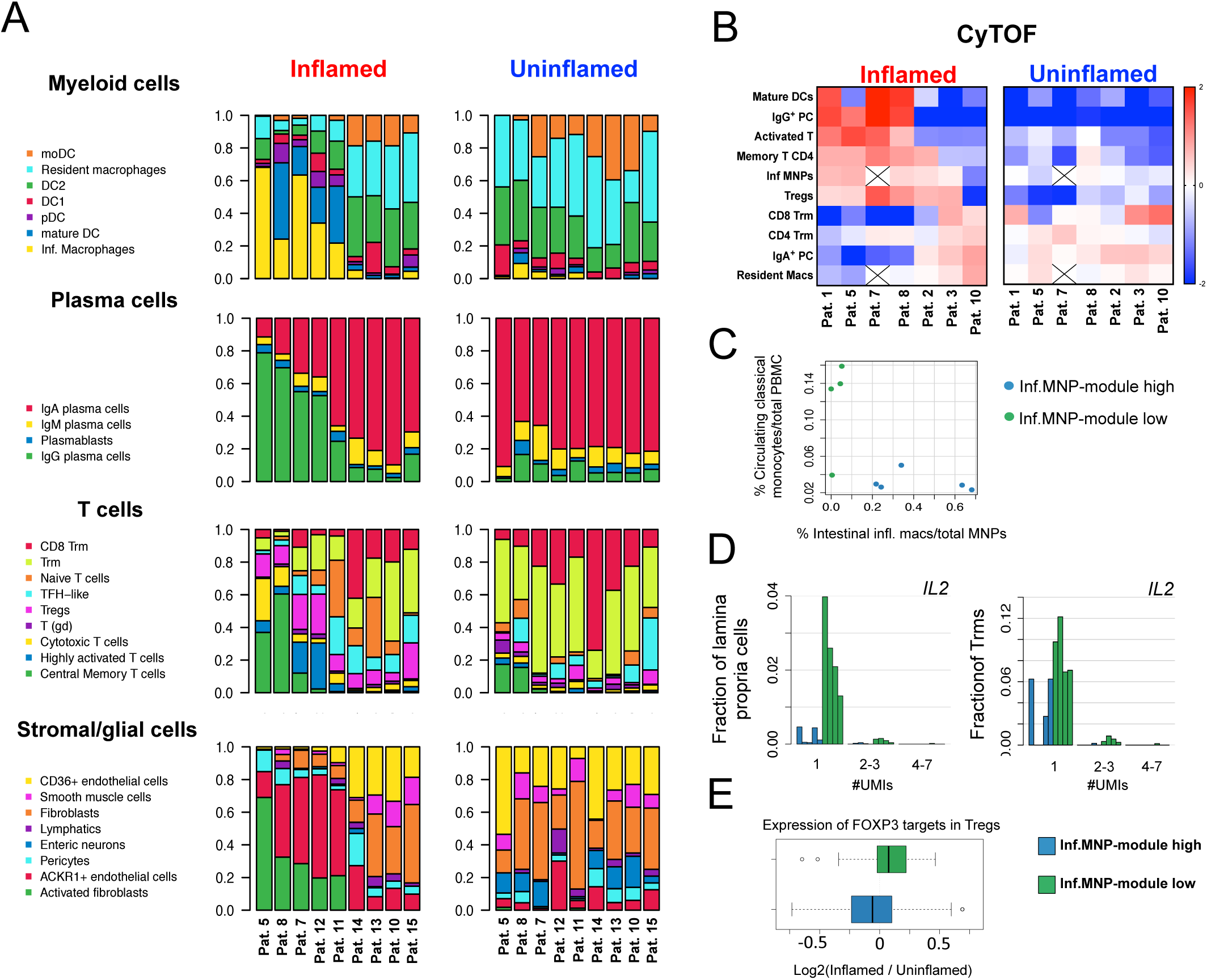
The Inf. MNP-module is enriched in a subset of ileal Crohn’s disease patients. **(A)** Analysis of cell subtype frequencies within their cellular compartment reveals prominent enrichment of the Inf.MNP-module in a subgroup of patients. Cellular composition of four cellular compartments are shown separately in inflamed and uninflamed samples of each patient. Patients are ordered by hierarchical clustering of frequency data in inflamed ileum. **(B)** CyTOF data confirm prominent enrichment of the Inf.MNP-module in a subgroup of patients. Heat-map showing the log2-normalized frequency of indicated cell subtypes (rows) in individual inflamed and uninflamed patient tissues (columns). **(C)** Scatter plot showing frequencies of blood classical monocytes (y-axis) vs. intestinal inflammatory macrophages (x-axis) in Inf.MNP-module high (blue dots) and Inf.MNP-module low (green dots) patients (**D**) Decreased *IL2* expression associates with the presence of the Inf-MNP-module. Histograms showing the fraction of total lamina propria cells (left) or resident memory T cells (right) per expression bin (1, 2–3, 4–7 UMIs) per Inf.MNP-module high (blue bars) or Inf.MNP-module low (green bars) patients. Cells were down-sampled to 1000 UMIs. Each bar represents a cell fraction of individual patient. (**E**) Box-plots showing the distribution of log ratios between average expression of FOXP3-target genes in regulatory T cells (Tregs) from inflamed vs. uninflamed ileum of Inf.MNP-module high and Inf.MNP-module low patients. The expression of FOXP3 targets is significantly lower in Tregs from inflamed ileums of Inf.MNP-module high patients (K.S. D=0.36; p=10^−5^).

### Enrichment for the Inf.MNP-module at diagnosis associates with resistance to anti-TNF

We next tested whether the Inf.MNP-module was also detectable in 4 independent cohorts of iCD patients (n=441). In line with previous cancer studies (Puram et al., 2017), we developed a straight-forward scoring function for normalized bulk-RNA samples to generate two gene expression scores accounting for the transcriptional variability between Inf.MNP-module high and Inf.MNP-module low patients in inflamed ileum. First, we defined gene lists associated with the relative abundance of the Inf.MNP-module through a differential expression analysis between pooled gene expression profiles of the two subgroups of patients identified by our scRNAseq analysis (**Figure 5A, Table S6**). Then, we projected bulk RNA microarray and sequencing data of endoscopic inflamed ileum pinch biopsies obtained from the 4 independent cohorts of iCD patients (**supplemental methods**) onto the two signatures by averaging the gene-lists z-scores for each sample (**Figure 5B**). For all 4 cohorts, a significant negative correlation existed between the 2 scores in the inflamed tissues of iCD patients, whereas no negative correlations could be detected in uninflamed ileums (**Figures 5C and S6A**), confirming the scRNA-seq results. Importantly, enrichment of the Inf.MNP-module was also detectable in early stages of the disease, prior to any biological therapy (RISK and UNITI-2; n=340 patients) (Feagan et al., 2016; Haberman et al., 2014; Kugathasan et al., 2017). The correlation between individual genes from each score was restricted to inflamed lesions (**Figure 5D, Figure S6B**), which indicated that distribution of the scores reflected the enrichment of the Inf.MNP-module in the inflamed ileums of a subset of patients in all 4 cohorts of iCD patients.

**Figure 5.**
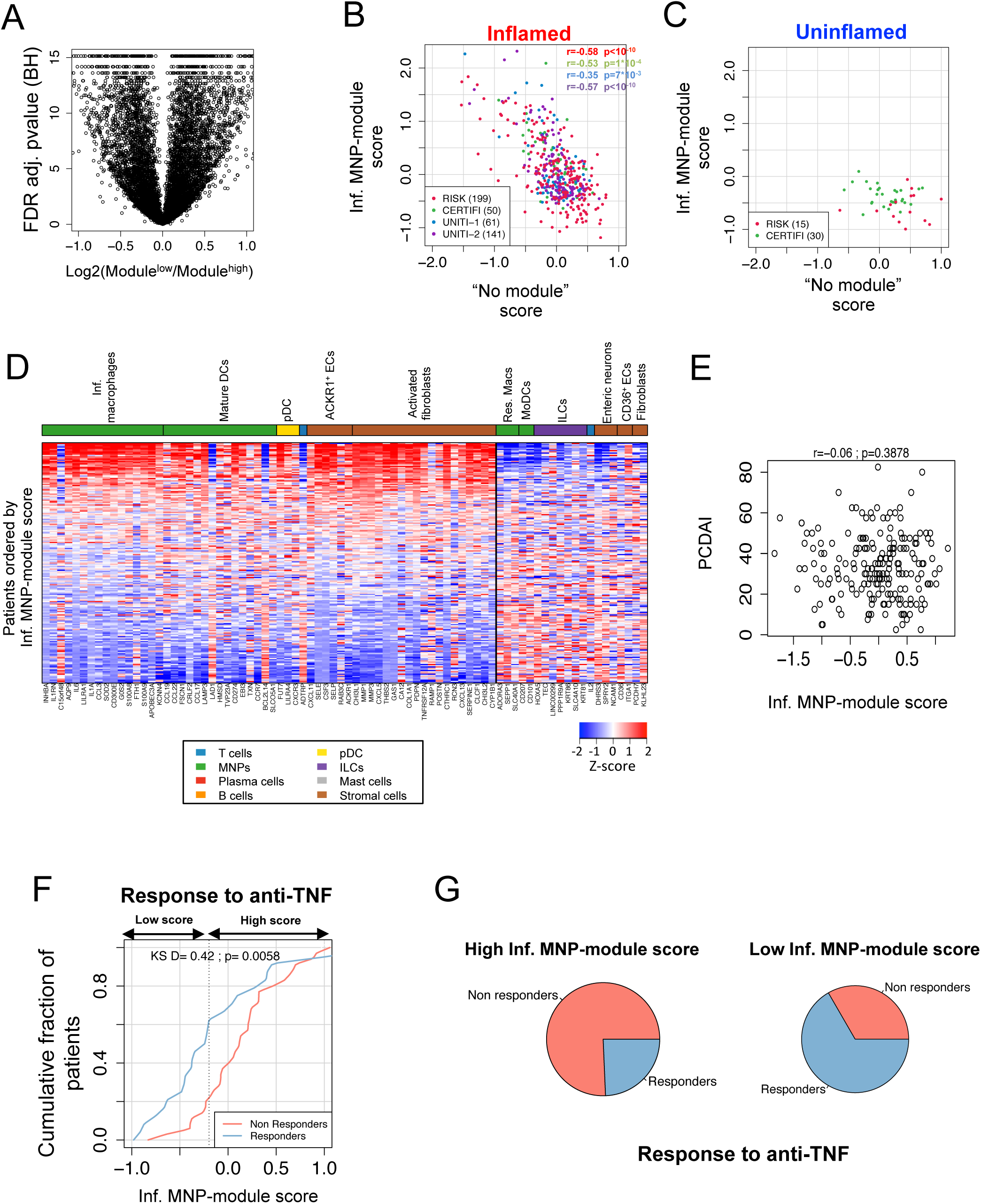
Enrichment for the Inf.MNP-module before treatment associates with resistance to anti-TNF. **(A)** Differential expression analysis between pooled scRNAseq data from inflamed samples of patients stratified by Inf.MNP-module enrichment. Volcano plot showing for each gene the log2 ratio between averages in Inf.MNP-module high and Inf.MNP-module low patients (x-axis), and minus Log2 BH adjusted p-value (y-axis). See methods for detailed description of the permutation test that provided the empirical p-values. **(B-C)** Validation of the Inf.MNP module by bulk RNA datasets. Scatter plots showing the projected bulk RNA microarray and sequencing data of inflamed (B) and uninflamed (C) biopsies from 4 Crohn’s disease cohorts onto the signature scores (see methods). The negative correlation over inflamed samples indicates the enrichment of the Inf.MNP-module in a subset of patients in each cohort. **(D)** Heat-map of normalized bulk RNAseq expression values (read-high; blue-low) of selected genes (columns) among CD inflamed ileal biopsies (stacked rows) ordered according to their enrichment for the Inf-MNP-module-high signature. (**E**) Pediatric Crohn’s disease index (PCDAI) does not correlate with the Inf.MNP-module score in newly diagnosed patients before anti-TNF therapy. Shown is PCDAI of RISK patient (y-axis) vs. Inf.MNP-module score at diagnosis (x-axis). Pearson correlation test (r=-0.06) was not significant. **(F)** High Inf.MNP-module score is associated with non-response to anti-TNF therapy. Cumulative distribution of Inf.MNP-module scores in responders and non-responders to anti-TNF at diagnosis before treatment. Non-responders have significantly higher Inf.MNP-module scores (K.S. D=0.42; p=0.006). Dashed line indicates the score value corresponding to K.S. D statistic (score=-0.2). **(G)** Pie charts showing the enrichment for responders and non-responders to anti-TNF in patients classified as Inf.MNP-module high or low according to K.S. D statistic.

The introduction of anti-TNF antibodies has transformed the clinical outcome of IBD patients but 20–30% of iCD patients never respond to these drugs, and continue to undergo uncontrolled bowel damage requiring surgical intervention within 10 years after the diagnosis (Torres et al., 2017). Interestingly, although expressed by infl. macs, TNF was not preferentially associated with the Inf.MNP-module, likely because of additional T cell sources in inflamed tissues (**Figures 3F&G**), and raising the possibility that patients enriched for the Inf.MNP-module could respond differently to TNF blockade. We next tested if enrichment for the Inf.MNP-module impacted the response to anti-TNF therapy in iCD patients. We focused our analysis on patients from the RISK cohort, which is a prospective inception cohort of pediatric patients with newly diagnosed Crohn’s disease (Kugathasan et al., 2017), for whom long-term clinical follow-up was available. We selected patients who received anti-TNF antibodies within their first year of disease and assessed clinical remission between 18 and 24 months after diagnosis, defining patients as responders only if they achieved durable corticosteroid-free clinical remission. The Inf.MNP-module score did not correlate with the pediatric Crohn’s disease activity index (PCDAI) at diagnosis (**Figure 5E**). Strikingly, however, the Inf.MNP-module score at baseline showed significantly distinct distributions between non-responder and responder populations (Kolmogrov-Smirnov D=0.42; p=0.006) (**Figure 5F)**, with a major enrichment of anti-TNF non-responders in patients with high Inf.MNP-module scores prior to initiation of anti-TNF treatment (**Figure 5G**). Similar to our discovery cohort, Inf.MNP-module high patients could not be distinguished by markers of systemic inflammation in the RISK cohort, suggesting that traditional CD endpoints were unable to predict disease response to biological therapy (**Figure S6C**).

Taken together, our data thus identified the presence of an Inf.MNP-module in inflamed tissues of a subgroup of iCD patients, which associated with primary non-response to anti-TNF therapy.

## Discussion

Pathophysiologic heterogeneity between patients has been proposed as a major cause for the failure of novel drug trials in IBD patients in the past two decades (Bilsborough et al., 2016). In this study, we used single cell technologies to characterize the ileal lamina propria cellular landscape in individual patients to an unprecedented level of granularity and identified a pathogenic Inf.MNP-module prominently enriched in lesional tissues from a subgroup of patients. We validated the molecular signature of the Inf.MNP-module in multiple bulk expression datasets and correlated the presence of the module with primary non-response to anti-TNF in a pediatric inception cohort (RISK).

The Inf.MNP-module includes two populations of pathogenic MNP, namely infl. macs and mature DCs. In line with previous studies, ligand/receptor analyses suggest that infl. macs contribute to stromal cell activation partly through the production of inflammatory cytokines IL-6 and IL-1β (Kamada et al., 2008; Schenk et al., 2007), and OSM, another IL-6 family member recently shown as a significant inducer of pathogenic stromal cell activation in IBD tissues (West et al., 2017). Intestinal macrophages in mice are heterogeneous and arise from the CCR2-dependent replenishment by blood classical monocytes (Bain et al., 2014) and from embryonic precursors that are recruited to the intestine prior birth (De Schepper et al., 2018). In the healthy intestine, tissue resident macrophages are epigenetically wired by local microenvironmental cues to acquire a regulatory program necessary for the establishment and preservation of intestinal homeostasis (Lavin et al., 2014; Mowat et al., 2017). Part of this program comprises the acquisition of inhibitory mechanisms limiting the production of inflammatory cytokines in response to inflammatory stimuli by gut-resident macrophages (Smythies et al., 2005, 2010). A previous work suggested decreased blood levels of classical monocytes in the blood of CD patients (Thiesen et al., 2014). We suggest that classical monocyte reduction preferentially occurs in the blood of Inf.MNP high patients and may be driven or maintained by activated stromal cells that are highly enriched in inflamed lesions of these patients. Activated stromal cells in inf.MNP lesions include activated fibroblasts which produce the CCR2 ligands CCL2 and CCL7, and activated endothelial cells that express the atypical chemokine receptor ACKR1, which binds tissue-derived CCL2 and CCL7 to facilitate monocyte transcytosis in inflamed tissues (Pruenster et al., 2009). Our data suggest a scenario in which dysregulated stromal cells in Inf.MNP-module high patients facilitate the infiltration of blood classical monocytes in inflamed ileum to generate pathogenic infl. macs, which in turn further amplify stromal cell activation. These results support and extend previous data identifying hemopoietic-derived OSM/stromal interactions as a central hub in IBD pathology (West et al., 2017). Interfering with the sustained feeding of the inflammatory macrophage pool by blood classical monocytes using anti-CCR2 blocking antibodies could help disrupt the inflammatory macrophages/stromal hub, while preserving tissue-protective gut-resident macrophages. Interestingly, activated fibroblasts also expressed IL-11, which beyond recently identified profibrotic properties, has been suggested to exert neuroprotective functions (Schafer et al., 2017; Zhang et al., 2011). Both Inf.MNP-module high and Inf.MNP-module low patients, however, exhibited similar reductions of enteric neurons in inflamed tissues, suggesting that fibroblasts-derived IL-11 was not enough to maintain neuronal integrity in inflamed ileums.

Infl.MNP-high patients are also enriched in mature DC, which in contrast to inflammatory macrophages have been poorly characterized in IBD lesions (Hart et al., 2005; Magnusson et al., 2016; Middel et al., 2006). We show that mature DC in the lamina propria of Inf.MNP lesions are located within lymphoid aggregates of T and B cells and express high levels of the chemokine receptor CCR7, high levels of the CCR7 ligand CCL19, as well as CCR4 ligands, indicating potential privileged and pathogenic interactions with CCR4/CCR7 expressing T_CM_. Concordant with the function of mature DC in T cell response initiation, mature DC expressed high levels of co-stimulatory molecules together with higher protein levels of HLA-DR at their membrane. These data strongly suggest that mature DC likely control the local recruitment, activation, expansion and spatial organization of a pathogenic adaptive immune response, supported by the presence of highly activated T cells, CTLs and IgG-plasma cells, which represent a second central hub in the formation of the Inf.MNP-module. In addition, we found that Tregs expressed lower FOXP3 and FOXP3 target genes suggestive of defective Tregs response in these patients.

Strikingly, while we first identified the pathogenic module in a small group of patients with advanced clinical and histological diseases, we confirmed the presence of the Inf. MNP module in 4 independent iCD cohorts (n=441) and showed that presence of this pathogenic module at diagnosis correlated with primary resistance to anti-TNF therapy.

Improper control of mucosal inflammation in ileum CD represents a significant risk factor of progressive bowel damage accumulation and surgical resection (Pariente et al., 2011). Here we suggest that an Inf.MNP-module centered around two major pathogenic hubs organized by mature DC and monocyte-derived inflammatory macrophages associates with resistance to anti-TNF. By providing a comprehensive cellular and molecular basis for the resistance to anti-TNF observed in a subgroup of patients (**Figure S7**), our study opens novel opportunities for therapeutic discoveries tailored to primary non-responders to anti-TNF.

By linking deep profiling by scRNAseq to identify complex cellular and molecular signatures in subgroup of patients with the breadth achieved by low-cost bulk RNA sequencing methods applied to hundreds of patients necessary for validation and clinical interpretation, our approach benefited from the power to connect high-level clinical parameters such as response to treatment, with specific molecular phenotypes (i.e. specific cellular subtypes, receptors/ligands), which can be tested in follow-up studies. This approach of combining high resolution single cell mapping of inflammatory lesions in small numbers of patients with bulk RNA sequencing on large cohorts where extensive clinical characterization is generalizable, highlights the potential to broadly transform understanding of human multifactorial immune-mediated inflammatory diseases.

## Acknowledgments

The authors would like to warmly thank Dr Jean-Frederique Colombel (Co-Director Feinstein IBD Center; Mount Sinai Hospital, New York, NY) for invaluable discussions and insights on Crohn’s disease. The authors would also like to thank Dr Helene Salmon (Precision Immunology Institute, Icahn School of Medicine at Mount Sinai, New York, NY) for careful reading of the manuscripts and precious discussions on stromal cells analysis. The authors thank Dr Emilie Grasset (Precision Immunology Institute, Icahn School of Medicine at Mount Sinai, New York, NY) for critical insights on B cells analysis. We thank Dr Andrew Chess for fruitful discussions. Funding: JCM is supported by “Prix pour les jeunes chercheurs” de la Fondation Bettencourt-Schueller and by the Philippe Foundation; RCU is supported by a Career Development Award from the Crohn’s and Colitis Foundation.

## Author contributions

Conceptualization, J.C.M., E.K., J.H.C., M.M.; Methodology, J.C.M; G.B., E.K., G.A., S.G, A.R.; Software, E.K.; Formal Analysis, E.K.; Investigation, J.C.M, G.B; C.C, M.G., L.W, A.L, L.W, H.M.K, I.L.; Data Curation, M.G, R.C.U; Resources, L.S.C; S.N., A.G., M.D., J.S.F, C.E.W, L.M., S.K., L.A.D, J.H.H, J.F., P.D., H.M.K, S.G., A.R.; Writing – Original Draft, J.C.M, E.K, M.M.; Writing – Review & Editing, C.W.E., J.H.C., M.M.; Visualization, J.C.M.; C.C., G.A., E.K.; Supervision, M.M., J.H.C., E.K.; Funding Acquisition, J.H.C., M.M.

---

**Figure S1.**
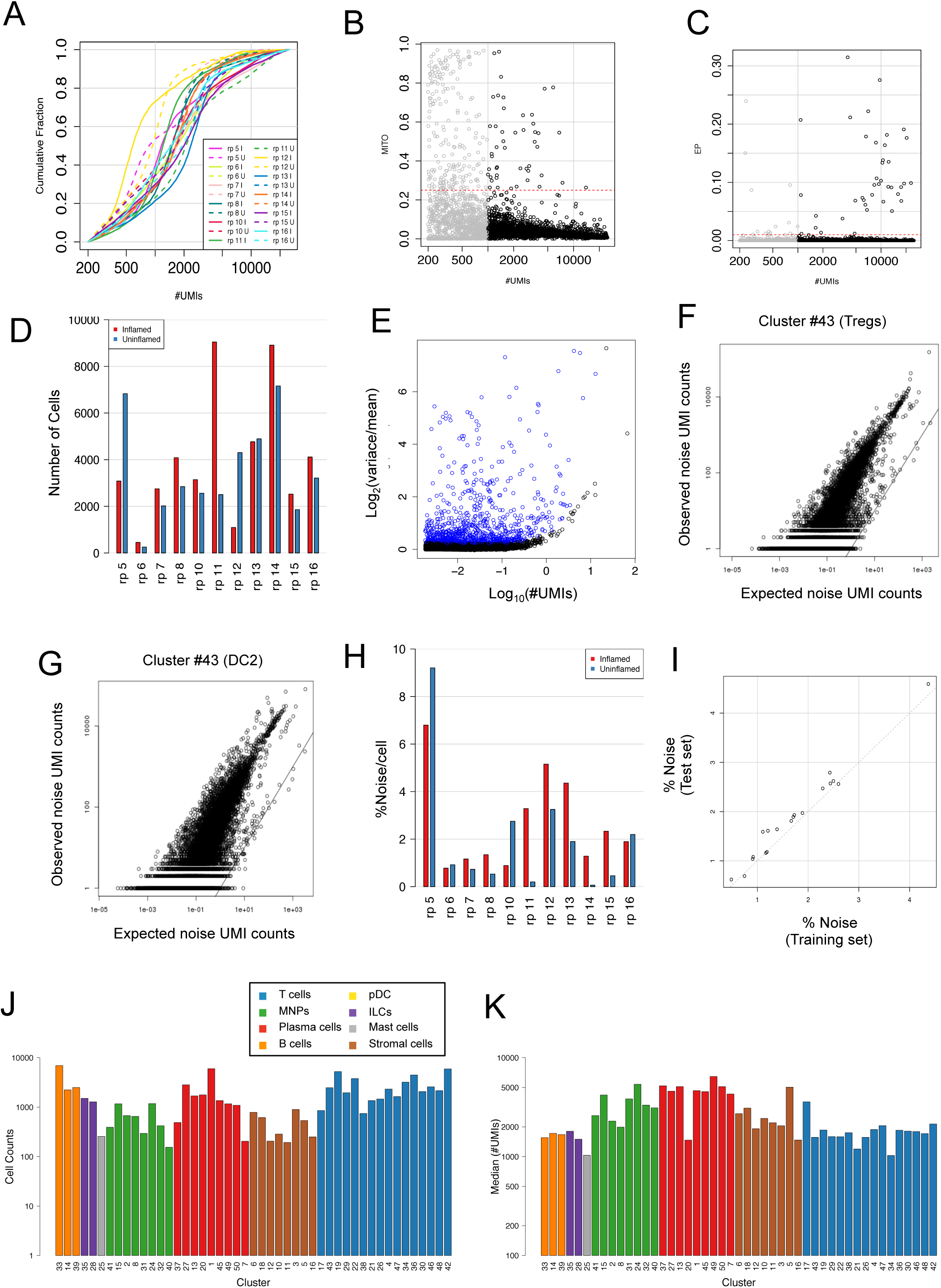
Clustering analysis of 22 inflamed and uninflamed ileum CD tissues, Related to Figure 1. **(A)** Cumulative fraction (y-axis) of UMI counts per cell (x-axis) for inflamed (solid lines) and uninflamed (dashed lines) samples **(B)** Filtering low-quality cell-barcodes. Shown are UMIs fraction of mitochondrial genes (y-axis) vs. the number of UMIs (x-axis) per cell-barcode for a representative sample (patient 7 inflamed ileum). Cell-barcodes below 1000 UMIs were filtered (gray dots) while cell-barcodes with mitochondrial fraction lower than 0.25 (red dashed line) were included for further analysis. **C)** Similar to B but showing filtering of epithelial cells. Cell-barcodes with more than 1% epithelial UMI fraction were filtered. **(D)** Number of cells per patient per tissue recovered after QC steps. **(E)** Selection of variable genes for clustering initiation. Shown is the variance divided by the mean vs. the mean for genes included in the initiation step of the clustering. UMI matrix was down-sampled to 557 UMIs. **(F-G)** Estimation of technical noise. Shown are observed UMI counts vs. the expected number of noise UMIs per gene for Treg (F) and DC2 (G) clusters with the identity line (solid). Genes that are usually not expressed in these cell subtypes, serving as endogenous “spike-ins” and almost all their UMIs in these clusters are predicted to be noise UMIs. **(H)** Estimated noise percentages per sample. **(I)** Cross validation of noise parameters. Noise percentage per sample as estimated for the test set (y-axis) vs. the training set (x-axis). **(J)** Number of cells per cluster. Clusters are grouped by cellular compartment. **(K)** Median number of UMIs/cells for each cluster.

**Figure S2.**
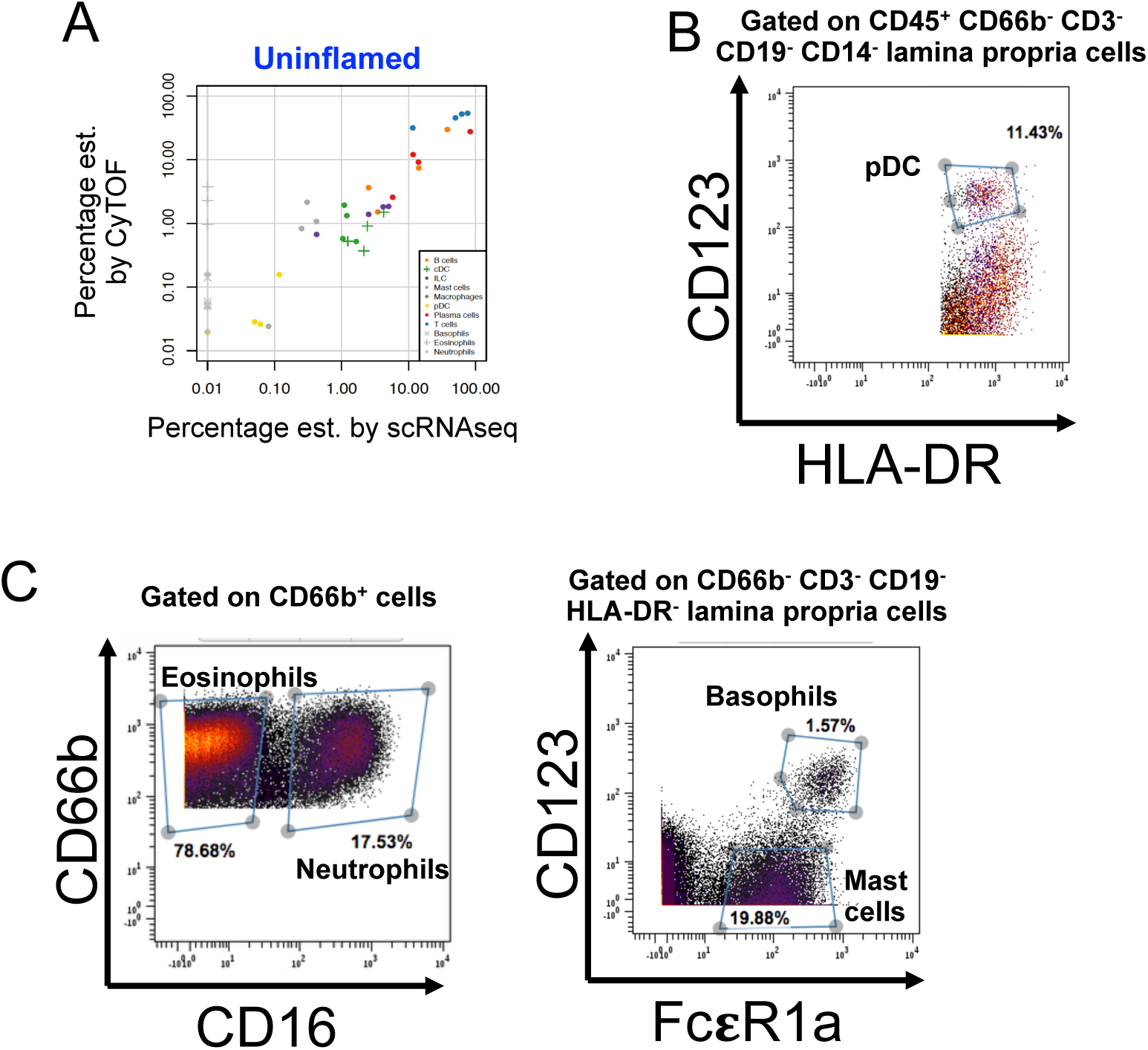
CyTOF analysis validates immune lineage frequencies estimated by scRNAseq and identifies granulocyte populations, Related to Figure 1. (**A**) Scatter plot showing lineage frequencies estimated based on CyTOF (y-axis) vs. scRNAseq (x-axis) data of uninflamed ileums of CD patients (n=4 patients). (**B**) Representative gating of CD45^+^ CD66b^−^CD3^−^CD19^−^CD14^−^HLA-DR^+^ CD123^+^ plasmacytoid dendritic cells (pDCs). **(C)** Representative gating strategies to identify granulocyte populations by CyTOF analysis of CD ileal tissues. Left: Representative gating of CD45^+^ CD66b^+^ CD16^+^ neutrophils and CD45^+^ CD66b^+^ CD16^−^eosinophils. Right: Representative gating of CD45^+^ CD66b^−^CD3^−^CD19^−^HLA-DR^−^ FcεR1a^+^ CD123^+^ basophils and CD123^−^mast cells.

**Figure S3.**
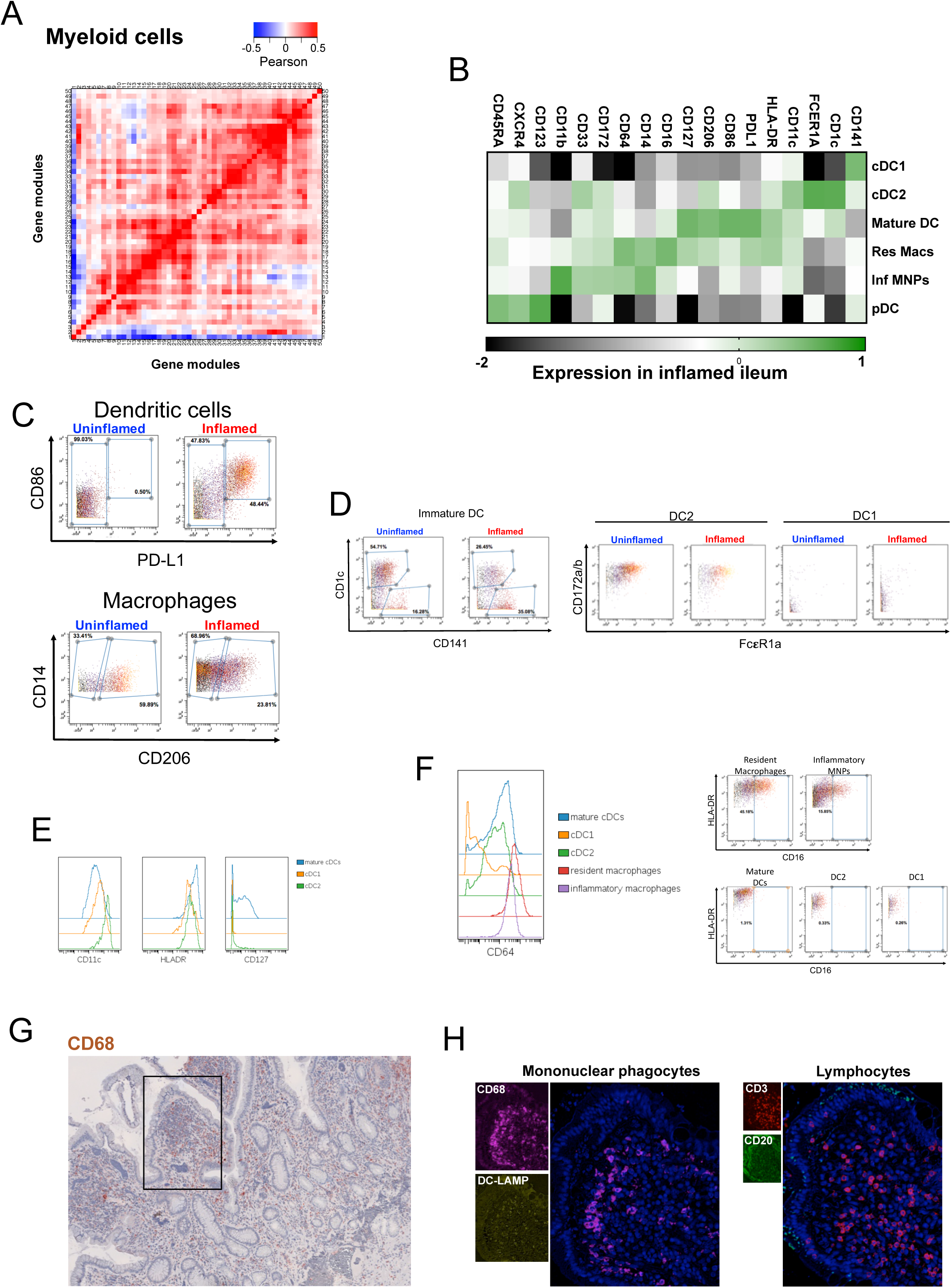
Validation of the myeloid cells diversity at the protein level, Related to Figure 2. **(A)** Correlation matrix of modules defined by genes with strong covariance of expression within the myeloid cells. **(B-F)** Validation of the scRNAseq analysis of MNP at the protein level. **(B)** Heat map showing the color-coded log-normalized expression of membrane markers between myeloid cell populations analyzed by CyTOF (n=7 CD patients). **(C)** Top: Representative gating of mature DC in uninflamed and inflamed ileum. Validating the scRNAseq clustering analysis (**Figure 2C**), mature DC were defined by the co-expression of CD86 and PD-L1 within CD45^+^ CD66b^−^CD3^−^CD19^−^HLA-DR^+^ CD14^−^CD16^−^CD123^−^CD11^high^ cells. Bottom: Representative gating of CD45^+^ CD66b^−^CD3^−^CD19^−^HLA-DR^+^ CD14^+^ macrophages to identify CD206^+^ resident and CD206^−^inflammatory macrophages as defined by the scRNAseq analysis (**Figure 2D**). **(D)** Left: Representative gating of immature DC in uninflamed and inflamed ileums to identify CD141^+^ DC1 and CD1c^+^ DC2. For DC1, CD141 was used as a surrogate of *CLEC9A* expression in the scRNAseq analysis, as previously reported (Villani et al., 2017). Right: Representative gating showing specific expression of FcεR1a (*FCER1A*) on DC2. **(E)** Histograms showing higher HLA-DR and lower CD11c levels on CD86^+^ PD-L1^+^ mature DC as compared to immature DC1 and DC2, concordant with the transcriptomic similarities between mature DC in the ileum CD lamina propria and migratory DC (Miller et al., 2012). Validation of the restricted expression of CD127 (*IL7R*) on mature DC as identified in the scRNAseq analysis. **(F)** Left: Histograms confirming higher CD64 (*FCGR1A*) expression in macrophages vs DC. Right: Representative staining validating the restricted expression of CD16 (*FCGR3A*) to macrophages. **(G)** Representative staining of CD68^+^ macrophages in the inflamed lamina propria. The black square identifies the region with a mature DC-centered lymphoid aggregate presented at higher magnification in main **Figure 2F**. **(H)** Representative staining of a resident macrophages-T cell aggregate devoid of mature DC and B cells.

**Figure S4.**
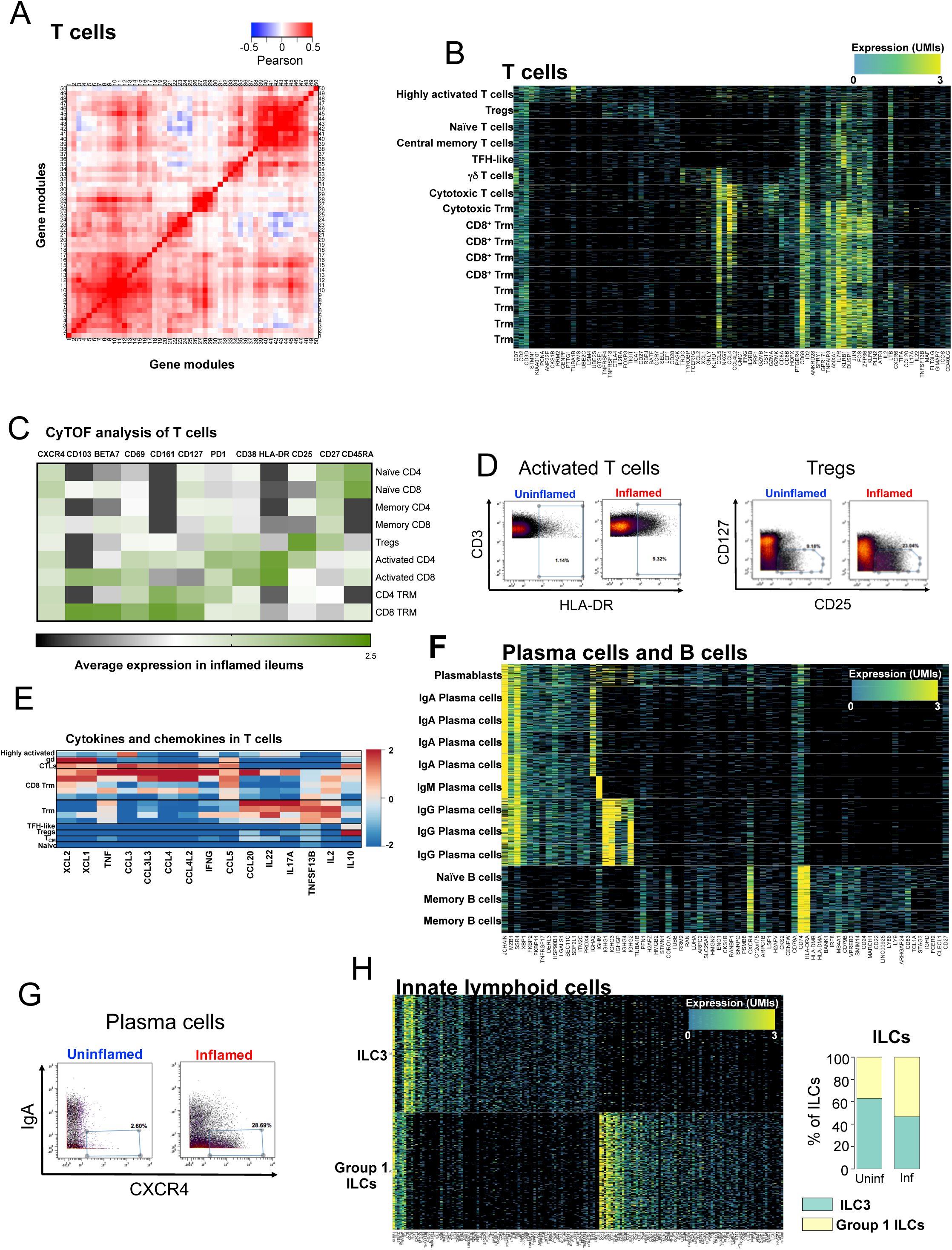
Characterization of lymphocyte populations iCD, Related to Figure 3. **(A)** Correlation matrix of modules defined by genes with strong covariance of expression within the lymphoid cells. **(B)** T cell clusters. Stacked columns correspond to single-cells and rows to selected differentially expressed genes for which down-sampled UMI counts are color-coded. Clusters are demarcated by gray bars. **(C)** Heat map showing the color-coded normalized expression of membrane markers between T cell populations analyzed by CyTOF (n=7 CD patients). **(D)** Left: Representative gating of HLA-DR^+^ activated T cells in uninflamed and inflamed ileum. Right: Representative gating of CD25^high^ CD127^low^ Tregs in uninflamed and inflamed ileum. **(E)** Heat map showing relative expression of cytokines and chemokines between T cell populations. **(F)** B cell clusters. Stacked columns correspond to single-cells and rows to selected differentially expressed genes for which down-sampled UMI counts are color-coded. Clusters are demarcated by gray bars. **(G)** Representative gating of IgA^−^CXCR4^+^ plasma cells corresponding to IgG producers, in uninflamed and inflamed ileum. **(H)** Innate lymphoid cell clusters. Left: Stacked columns correspond to single-cells and rows to selected differentially expressed genes for which down-sampled UMI counts are color-coded. Right: Relative enrichment of ILC1/NK cells and ILC3 in uninflamed and inflamed ileums.

**Figure S5.**
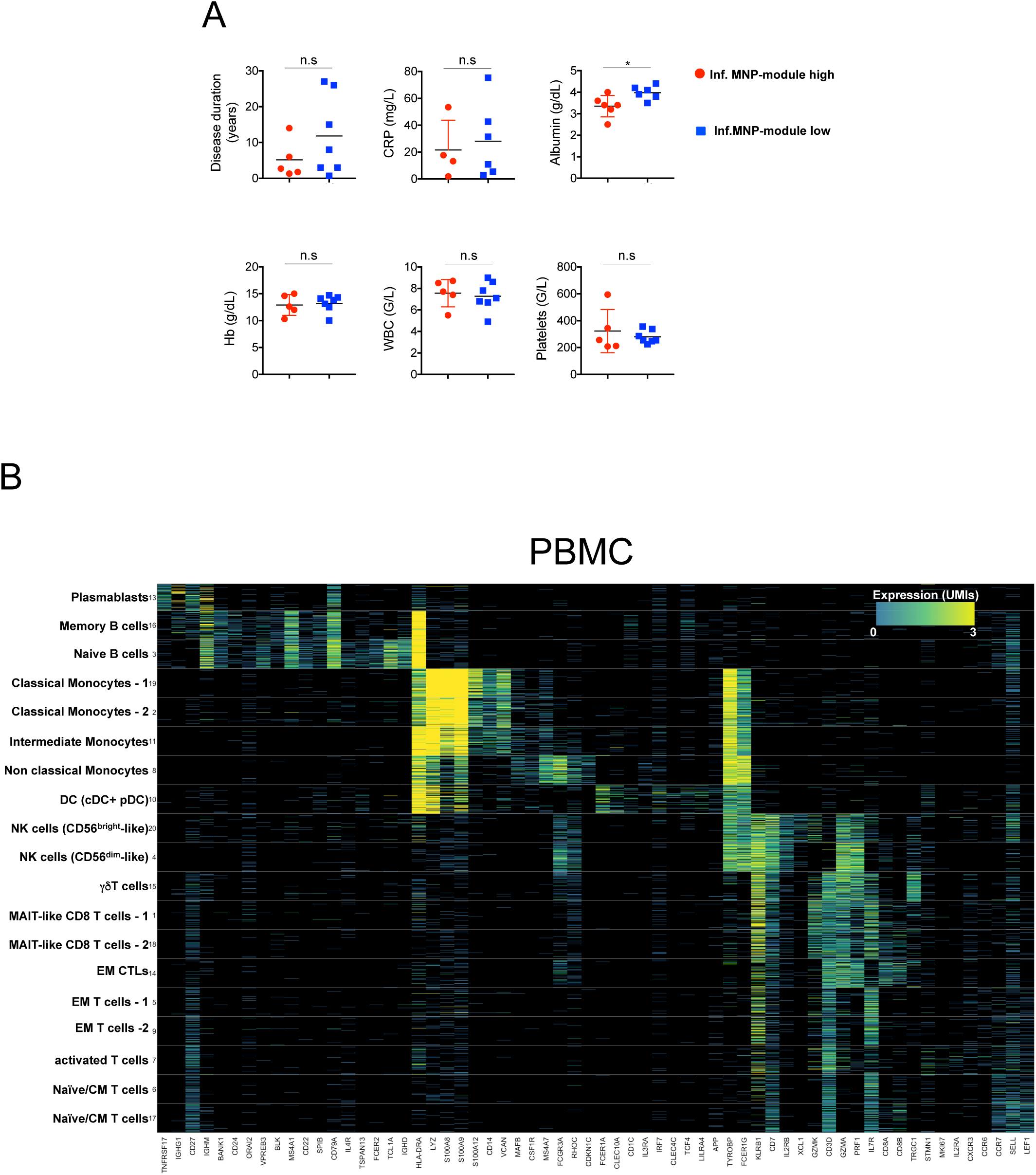
Similar distributions of clinical parameters in Inf.MNP-high and Inf.MNP-low patients, Related to Figure 4. **(A)** Comparison of clinical parameters between Inf.MNP-module high and Inf.MNP-module low patients. Patients analyzed by scRNAseq and/or CyTOF are included in the comparison. CRP: C-reactive protein; WBC: white blood count; ESR: erythrocyte sedimentation rate; Hb: hemoglobin; Alb: albumin. (**B**) PBMC clustering analysis. Stacked columns correspond to single-cells and rows to selected differentially expressed genes for which down-sampled UMI counts are color-coded. Clusters are demarcated by gray bars.

**Figure S6.**
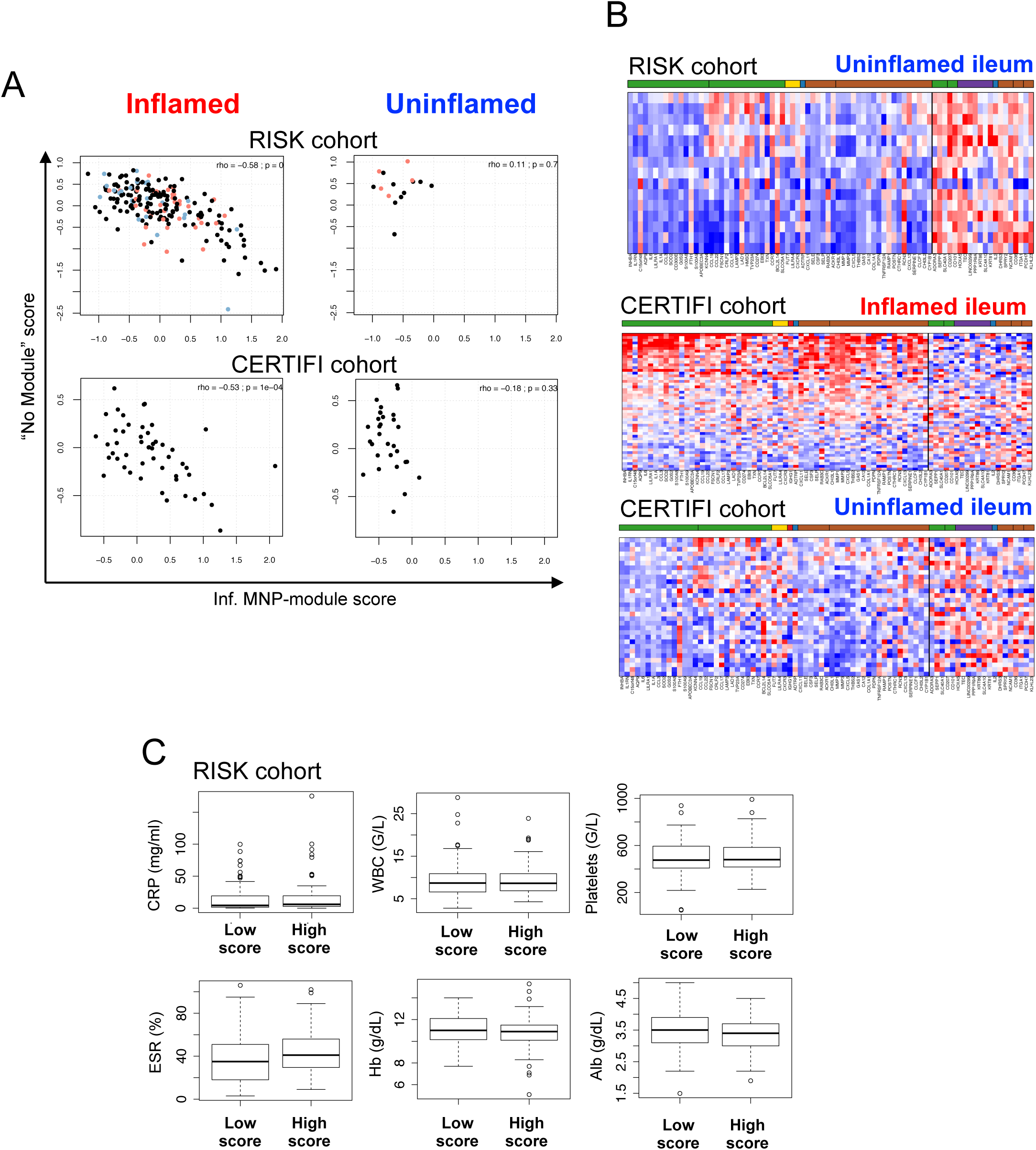
Blood markers of inflammation don’t predict Inf.MNP-module high scores, Related to Figure 5. **(A)** Scatter plot showing the projected bulk RNA microarray and sequencing data of inflamed biopsies from the RISK (top) and CERTIFI (bottom) cohorts onto the signature scores defined after the scRNAseq analyses to detect enrichment of the Inf.MNP-module. **(B)** Heatmap of normalized bulk RNA-seq gene expression values (read-high; blue-low) for selected genes (columns) for different CD patients’ ileal samples (stacked rows), ordered according to their enrichment for the Inf-MNP-module signature score. Shown are uninflamed ileum from the RISK cohort, inflamed and uninflamed ileum from the CERTIFI cohort. **(C)** Whisker plots comparing markers of systemic inflammation in Inf.MNP-module high and Inf.MNP-module low patients from the RISK before anti-TNF therapy. CRP: C-reactive protein; WBC: white blood count; ESR: erythrocyte sedimentation rate; Hb: hemoglobin; Alb: albumin.

**Figure S7.**
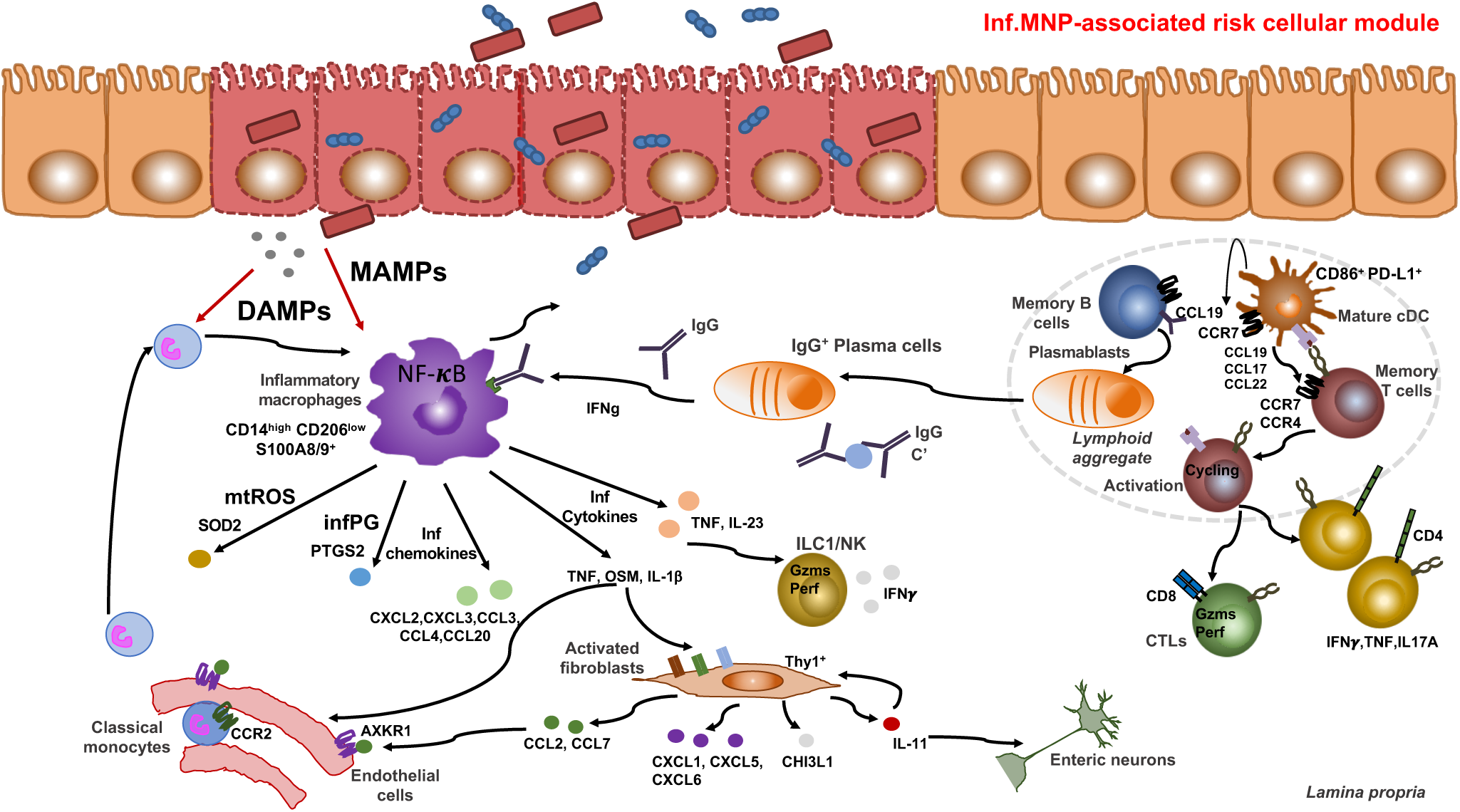
Graphical Summary. Graphical summary proposing a pathophysiological model of the Inf.MNP-associated immune response in the inflamed ileum of a subgroup of CD patients. DAMPs: danger associated molecular patterns. MAMPs: Microbial associated molecular patterns. cDC: conventional dendritic cells.

**Table S1.** Clinical characteristics of Crohn’s disease patients included for scRNAseq and CyTOF studies.

| **Characteristic** | **Patients (n=14)** |
| --- | --- |
| **General data** |  |
| Age (y.o.; mean ± sd) | 32.7 ± 12.9 |
| Sex (M/F) | 10/4 |
| Disease duration (y.; mean ± sd) | 9.1 ± 9.3 |
| Smokers – no. (%) | 1 (7) |
| **Montreal classification** |  |
| **Age at diagnosis – no. (%)** |  |
| A1 below 16 y.o. | 3 (21) |
| A2 between 17 and 40 y.o. | 10 (71) |
| A3 above 40 y.o. | 1 (7) |
| **Location – no. (%)** |  |
| L1 ileal | 13 (93) |
| L2 colonic | 0 (0) |
| L3 ileocolonic | 1 (8) |
| L4 isolated upper disease | 0 (0) |
| **Behavior – no. (%)** |  |
| B1 non-stricturing, non-penetrating | 0 (0) |
| B2 stricturing | 12 (86) |
| B3 penetrating | 2 (15) |
| **Ileal resection size** |  |
| Length (cm; mean ± sd) | 21.5 ± 10.5 |
| Circumference (cm; mean ± sd) | 4.9 ± 1.1 |
| **Pathology – no. (%)** |  |
| Active ileitis with transmural chronic inflammation | 13 (100) |
| Granuloma | 3 (21) |
| **Lab data** |  |
| Hemoglobin (g/dL; mean ± sd) | 13.2 ± 1.6 |
| Platelets (G/L; mean ± sd) | 286 ± 102 |
| Albumin (g/dL; mean ± sd) | 3.7 ± 0.5 |
| CRP (mg/L; mean ± sd) | 25.5 ± 24.7 |
| **Medication at time of surgery – no. (%)** |  |
| Antibiotics | 4 (29) |
| 5-ASA | 6 (36) |
| 6-mercaptopurine | 4 (29) |
| Corticosteroids | 7 (50) |
| Anti-TNFα | 10 (71) |

**Table S2. Metadata and technical statistics per sample**. Relates sample IDs with patient IDs, technical, experimental and sequencing information, as well as single-cell UMI statistics.

**Table S3. Cell counts per cluster per sample**. Indicates the number of cells in each cluster for each sample.

**Table S4. MNP-based gene modules**. Gene modules derived from gene-to-gene correlation analysis of MNP cells.

**Table S5. Differential expression analysis of Tregs between Inf.MNP-module high and Inf.MNP-module low patients**. Statistics per gene participating in the differential expression analysis of Tregs.

**Table S6. Differential expression analysis of all cells between Inf.MNP-module high and Inf.MNP-module low patients**. Statistics per genes participating in the differential expression analysis between cells from inflamed samples, regardless of clusters.

**Table S7. Average gene expression per subtype**. Total UMI counts per gene per subtype divided by total UMI counts per subtype. Values are log2 transformed.

**Table S8. T cells-based gene modules**. Gene modules derived from gene-to-gene correlation analysis of T cells.

## Materials and Methods

### Human specimens

Patients eligible for inclusion in the study were identified by screening surgical programs at the Mount Sinai Hospital. Ileum CD tissues and venous blood were obtained after 1^st^ surgical resection. Protocols were reviewed and approved by the Institutional Review Board (IRB) at the Icahn School of Medicine at Mount Sinai (Mechanisms of intestinal inflammation following Ileal resection for Crohn’s Disease; HSM#13–00998). Clinical characteristics, including Montreal classification, of the patients retained for scRNAseq and CyTOF studies are summarized in **Table S1**. All macroscopically inflamed tissues included in the study were confirmed by pathological examination as active ileitis with transmural chronic inflammation.

### Preparation of lamina propria single cell suspensions

Tissues were collected in ice cold RPMI 1640 (Corning Inc., Corning, NY) and processed within one hour after termination of the surgery. To limit biased enrichment of specific immune and stromal populations related to local variations in the intestinal micro-organization (Mowat and Agace, 2014), we pooled fifteen to twenty mucosal biopsies sampled all along the resected specimens using a biopsy forceps (EndoChoice, Alpharetta, GA) to prepare cell suspensions. Epithelial cells were dissociated by incubating the biopsies in an EDTA-enriched dissociation medium (HBSS w/o Ca^2+^ Mg^2+^ (Life Technologies, Carlsbad, CA) - HEPES 10mM (Life Technologies) - EDTA 5mM (Life Technologies)) at +37°C under 100 rpm agitation for two cycles of 15 minutes. After each cycle, biopsies were hand-shaken for 30s and then vortexed vigorously for another 30s. Biopsies were then washed in complete RPMI media previously put at RT, and transferred in digestion medium (HBSS with Ca^2+^ Mg^2+^ - FCS 2% - DNase I 0.5mg/ml (Sigma-Aldrich, St. Louis, MO) - Collagenase IV 0.5mg/ml (Sigma-Aldrich)) for 40 minutes at +37°C under 100 rpm agitation. After digestion, the cell suspension was filtered through a 70μm cell strainer, washed in DBPS/ 2% FCS/ 1mM EDTA and spun down at 400g for 10 minutes. After red blood cell lysis (BioLegend, San Diego, CA), cells were washed as before. Dead cells were depleted from the suspension using the dead cell depletion kit (Miltenyi Biotec, Bergisch Gladbach, Germany), following manufacturer’s recommendations. Viability of the final cell suspension was calculated using a hemocytometer and Trypan blue (Corning) exclusion and was routinely > 85%.

### Preparation of PBMC

Venous blood was collected intraoperatively in EDTA BD Vacutainer^®^ blood collection tubes (Becton Dickinson, Franklin Lakes, NJ). Blood was diluted in sterile PBS (Sigma-Aldrich), layered above a Ficoll-Paque PLUS density gradient media (GE Healthcare, Little Chalfont, UK) and centrifuged 20 minutes at 800g, +20°C, brakes off. PBMC were collected at the interphase and washed. After RBC lysis, dead cells were removed from the suspension using the dead cell depletion kit (Miltenyi Biotec). Viability of the final cell suspension was calculated as above and was routinely > 90%.

### CyTOF sample preparation and analysis

Tissue cell suspensions were first incubated with Rh103 intercalator (Fluidigm, San Francisco, CA) for 20 minutes at 37°C to label non-viable cells, and then washed and labeled with a panel of metal-labeled antibodies for 30 mins on ice. The samples were then fixed and permeabilized with BD Cytofix/Cytoperm (BD Biosciences) and incubated in 0.125nM Ir intercalator (Fluidigm) diluted in PBS containing 2% formaldehyde for 30 mins. The samples were then washed and stored in PBS containing 0.2% BSA at 4oC until acquisition. Immediately prior to acquisition, samples were washed once with PBS, once with de-ionized water and then resuspended at a concentration of 1 million cells/ml in deionized water containing a 1:20 dilution of EQ 4 Element Beads (Fluidigm). The samples were acquired on a CyTOF2 (Fluidigm) equipped with a SuperSampler fluidics system (Victorian Airships) at an event rate of <500 events/second. FCS files were manually pre-gated on Ir193 DNA^+^ events, excluding dead cells, doublets, and DNA^−^ negative debris. Samples were analyzed using Cytobank (Mountain View, CA).

### Antibodies used in the CyTOF panel

| Tag | Marker | Clone | Source |
| --- | --- | --- | --- |
| 89Y | CD45 | HI30 | Fluidigm |
| 113In | CD57 | HCD57 | Biolegend |
| 115In | HLA-ABC | W6/32 | Biolegend |
| 141Pr | CD326 | Ep-CAM | Biolegend |
| 142Nd | CD19 | HIB19 | Biolegend |
| 143Nd | CD45RA | HI100 | Biolegend |
| 144Nd | CD141 | M80 | Biolegend |
| 145Nd | CD4 | RPA-T4 | Biolegend |
| 146Nd | CD8a | RPA-T8 | Biolegend |
| 147Sm | IgA | 9H9H11 | Biolegend |
| 148Nd | CD16 | 3G8 | Biolegend |
| 149Sm | CD127 | A019D5 | Biolegend |
| 150Nd | CD1c | L161 | Biolegend |
| 151Eu | CD123 | 6H6 | Biolegend |
| 152Sm | CD66b | G10F5 | Biolegend |
| 153Eu | PD-1 | EH12.2H7 | Biolegend |
| 154Sm | CD86 | IT2.2 | Biolegend |
| 155Gd | CD27 | O323 | Biolegend |
| 156Gd | CXCR3 | G025H7 | Biolegend |
| 158Gd | CD33 | WM53 | Biolegend |
| 159Tb | CD103 | Ber-Act8 | Biolegend |
| 160Gd | CD14 | M5E2 | Biolegend |
| 161Dy | CD56 | B159 | Biolegend |
| 162Dy | CD64 | 10.1 | Biolegend |
| 163Dy | CD172a/b | SIRPa/b | Biolegend |
| 164Dy | CD69 | FN50 | Biolegend |
| 149Sm | FcεRIα | AER-37 (CRA-1) | Biolegend |
| 166Er | CD25 | M-A251 | Biolegend |
| 167Er | CD11c | Bu15 | Biolegend |
| 168Er | CD3 | UCHT1 | Biolegend |
| 169Tm | integrin b7 | FIB504 | Biolegend |
| 170Er | CD38 | HB-7 | Biolegend |
| 164Dy | CD161 | HP-3G10 | Fluidigm |
| 172Yb | CD206 | 15-2 | Biolegend |
| 154Sm | CXCR4 | 12G8 | Biolegend |
| 174Yb | HLADR | L243 | Biolegend |
| 175Lu | PD-L1 | 29E.2A3 | Fluidigm |
| 176Yb | CD54 | HCD54 | Biolegend |
| 209Bi | CD11b | ICRF44 | Fluidigm |

### Multiplexed immunohistochemical consecutive staining on single slide (MICSSS)

Four μm-thick formalin-fixed paraffin-embedded (FFPE) tissue sections on glass slides were backed at 37°C overnight, deparaffinized in xylene, and rehydrated in decreasing concentrations of ethanol. Then, tissue sections were incubated in retrieval solution (Citrate buffer, pH6) for antigen retrieval at 95°C for 30 minutes. Tissue sections were incubated in 3% hydrogen peroxide and in serum-free protein block solution (Dako, X0909) before adding the primary antibody for 1 hour at room temperature. After signal amplification using biotinylated-secondary antibody and streptavidin-horseradish peroxidase, chromogenic revelation was performed using 3-amino-9-ethylcarbazole (AEC). Slides were counterstained with hematoxylin, mounted with a glycerol-based mounting medium and scanned for digital imaging (Hamamatsu Nanozoomer S60 Digital Slide Scanner). Then the same slides were successively stained as described (Remark et al., 2016). Primary antibodies were: anti-human CD3 (clone 2GV6, Ventana), anti-human CD20 (clone L26, Dako), anti-human DC-LAMP (clone 1010E1.01, Novus biologicals), anti-human CD68 (clone KP1, Dako) anti-human FOXP3 (clone 236A/E7, Abcam), anti-human CD206 (polyclonal, Abcam), and anti-human HLA-DR (clone TAL1B5, Abcam), anti-Ki67 (clone 30–09, Ventana).

### Processing of MICSSS pictures

Images were processed by an experienced pathologist. Whole slide images obtained from scanned immunostains performed on the same tissue sections were opened simultaneously on our digital pathology image analysis software (QuPath 0.1.3). Regions of interest (ROI) were defined using the same coordinates for each image. Images were exported as .png formatted images for analysis with the Fiji platform (Fiji - ImageJ 1.52h). Images are aligned by using the Trakem2 plugin. For each image, color deconvolution was achieved with a H-DAB vector to split images into three 8-bit channels including hematoxylin (blue), DAB (brown) and residual (green) channels. One hematoxylin channel was selected to define a base nuclear channel. All DAB channels were assigned different colors by using the Fiji look up tables (LUT) function. Colored channels were inverted and merged as one composite pseudo-fluorescence image. Brightness and contrast settings were optimized for each channel by comparing with original images. Underlying image pixel values were not changed for brightness and contrast settings.

### Molecular profiling

Patients 5–16 were analyzed by scRNAseq (**Table S2**). As discussed below, in order to minimize technical biases, the cell subtype clustering model was derived from scRNAseq samples acquired with the Chromium V2 chemistry. Cells from Patient 5, which were processed using the V1 chemistry, were projected onto the clustering model. Low cell number recovery prevented accurate high-resolution estimation of the cellular composition of Patient 6, who was consequently excluded from analyses involving correlations and comparisons of cell subtype frequencies. Because we did not observe significant differences between uninflamed and inflamed scRNAseq data from Patient 16, we also excluded this patient from analyses involving correlations and comparisons cell subtype frequencies.

### Droplet-based scRNAseq

Cells were suspended at 1.10^6^/ml in PBS and 10,000 cells were loaded onto the Chromium^TM^ Controller instrument within 15 min after completion of the cell suspension preparation using GemCode Gel Bead and Chip, all from 10x Genomics (Pleasanton, CA), and following the manufacturer’s recommendations. Briefly, cells were partitioned into Gel Beads in Emulsion in the Chromium™ Controller instrument where cell lysis and barcoded reverse transcription of RNA occurred. Libraries were prepared using 10x Genomics Library Kits and sequenced on an Illumina NextSeq500 according the manufacturer’s recommendations. Read-depth of more than 100 million reads per library, or an approximate average of 30,000 reads per cell was obtained (**Table S2**).

### Generation of gene expression matrices

FASTQ were demultiplexed using Cell Ranger v2.0 and aligned to the Grch38 human reference genome. Cell barcodes and unique molecular identifiers (UMIs) were extracted and “Raw” UMI matrix generated for each sample, providing with the number of UMIs for each gene that are associated with individual cell barcode. Cells enriched with mitochondrial, epithelial, or hemoglobin mRNA were excluded (see below **filtering low-quality cells and contaminating cell**).

### Filtering low-quality cells and contaminating cells

We extracted cell-barcodes associated with at least 800 UMIs from the “Raw” output UMI matrices of CellRanger (**Figure S1A)**, and filtered out cell-barcodes with at least 0.25% of mitochondrial mRNA (**Figure S1B**). Because collagenase digestion severely alters epithelial cell integrity, we also filtered out epithelial cells by excluding cell-barcodes with more than 1% UMIs associated with genes from the list below (**Figure S1B**). Potential residual red-blood cells were filtered out by excluding cell-barcodes with more than 10% UMIs associated with hemoglobin genes. Cell filtering statistics for the different samples are summarized in **Table S2**. Because of low cell counts recovery which prevented an accurate estimation of the frequencies of rare cellular subtypes, Patient 6 was excluded from analyses conducted after the clustering. The variability in cell-counts or UMI counts between the other samples was not confounding downstream analyses.

Epithelial gene list: PLA2G2A, CLCA1, REG4, S100A14, ITLN1, ELF3, PIGR, EPCAM, REG1B, REG1A, REG3A, FABP1, RBP2, SST, FABP2, SPINK1, FABP6, AGR2, AGR3, CLDN3, CLDN4, DEFA6, DEFA5, SPINK4, ALDOB, LCN2, MUC2, KRT8, KRT18, TSPAN8, OLFM4, GPX2, IFI27, PHGR1, MT1G, CLDN7, KRT19, FXYD3, LGALS4, FCGBP, TFF3, TFF1

### A batch-aware, multinomial mixture model for single-cell RNA-seq data

Analysis of background noise showed that noise profiles were proportional to the average gene expression profile of the sample as well as to the number of UMIs in the receiving cell. Our previous studies indicated that this type of cell-to-cell contamination could be attributed to molecular switching events that could occur during the amplification of the pooled cDNA library. We updated our previously published batch-aware mixture model (Jaitin et al., 2014; Paul et al., 2015) in order to accommodate the noise distribution observed here and defined the probability of observing gene i in cell j in the revised model as:

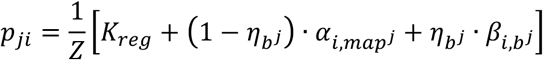

Where *map^j^* and *b^j^* are assignments of cells j to cell-type and batch(sample) respectively; 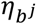 is the fraction of UMIs contributed by background noise; 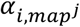 is the probability that a molecule drawn from celltype *map^j^* is of gene i (assuming no background noise) and 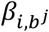 is the probability that a noise UMI drawn from batch *b^j^* will be of gene i.

Given this model and assuming hard association of cells with types, we can compute the log likelihood of the entire dataset as:

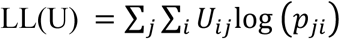

Based on the analysis of background noise distribution, we estimated the noise multinomial parameters by the average expression of the batch including all cell barcodes before filtering:

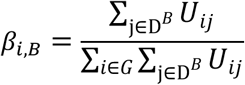

where D^B^ is a set of all barcodes with more than 100 UMIs in batch B and G is a set of all genes.

### Inference of the model parameters

We revised our previously described pseudo expectation-maximization (EM) algorithm to infer the assignment of cells to clusters *map^j^*, gene probability within cluster 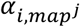 and background noise fraction 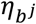. We implemented a cross-validation approach to test the robustness of the model’ parameters at each EM iteration and avoid over-fitting of the data. The updated algorithm outline was as follows:

1. Sample the single-cell sets:

A. Randomly sample without replacement 1000 cells from each batch to the learning set and 1000 cells to the test set. The resulting UMI matrices would be U^L^ and U^T^ respectively.
2. Initialize the model: Repeat A-F 10000 times

A. Randomly select a value N_ds_umis_ from the (P_1_P_2_) percentiles of the empirical distribution of the number of UMIs/cell of U^L^
B. Downsample U^L^ to N_ds___umis_ UMIs/cell. The downsampled matrix is denoted as U’^L^.
C. Select highly variable genes (see below)
D. Cluster the cells in U’^L^ based on the genes selected in step C using kmeans++ (https://tanaylab.bitbucket.io/tglkmeans/), with k seeds, following log2(X+K_reg___ds_) transformation.
E. Estimate *α, η* given kmeans++ assignments *map* for the cells in U’^L^ (see description below).
F. Given the current values of *α, η* parameters calculate the MAP assignment for each cell in U^L^ and update the assignments of cells to clusters *map^j^*,
G. Compute the log-likelihood of U^L^ to each of the current initialized types.
H. Select model parameters that correspond to the randomized seed that maximized the log-likelihood of U^L^
3. Estimate *α, η* given *map* for the cells in U^L^.
4. Given the values of *α, η* calculate the MAP assignment for each cell in U^L^ and update the assignments of cells to clusters map.
5. Return to step 3 and repeat until the likelihood converges, or the maximum number of iterations is reached.
6. Cross validation:

A. Given the values of *α, η* calculate the MAP assignment for each cell in U^T^ and update the assignments of cells to clusters *map*.
B. Estimate *α^T^, η^T^* given *map* for the cells in U^T^ and compare with *α, η*.
7. Estimate *η*, *map* given *α* for U

#### Selecting highly variable genes

Similarly as in other publications (Baran et al., 2018; Jaitin et al., 2014; Paul et al., 2015), we selected genes with variability that could not be explained by multinomial sampling variance. We calculated a loess curve for the log(variance/mean) vs. log(mean) distribution and binned the log(variance/mean) values by intervals of 0.2 of log(mean). We selected genes with more than 50 UMIs in U’^L^ from the 8^th^ percentile of each bin and also required that their log(variance/mean) is 0.1 or higher above the loess curve (**Figure S1E**).

#### Estimation of the model parameters

A. Estimate the model parameters. Repeat i-ii until convergence:

i. Subtract noise contribution from multinomial parameters (**Figures S1F-G**):

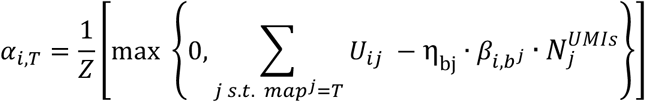

where 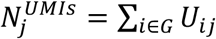; *G* contains all genes. Z is a normalization factor.
ii. Estimate the noise parameters **(Figures S1H-I**) by iterating over each batch b and selecting η_b_ that will optimize the likelihood of the cells in the batch 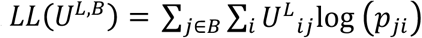 given constant *α* and *map*. Optimization process is simple since the likelihood components are additive.

For the joint clustering of the inflamed and uninflamed ileum samples we included barcodes with more than 1000 UMIs and used *K_reg_ds_* = 0.1; (P_1_,P_2_) = (10^th^,40^th^) percentiles; *K_reg_* = 5 · 10^−6^; k=50.

In the clustering analysis of PBMC samples we included all barcodes with more than 600 UMIs and used *K_reg___ds_* = 0.1; (P_1_,P_2_) = (10^th^,50^th^) percentiles; *K_reg_* = 5 · 10^−6^; k=20.

Permissive and fixed cutoffs on the number of UMIs were selected to allow subtypes with low UMI counts (**Figure S1K**) such as mast cells and specific subtypes of T cells to be included in the analysis across all samples. Findings of the study were not affected by modification of these thresholds within a range of 500–2000 UMIs/cells.

In order to improve the initiation of the model, genes with lateral expression across different subtypes were excluded from the k-means clustering (step 2.D.). These gene consisted of the following groups: cell-cycle, stress, ribosomal and mitochondrial, metallothionein genes, immunoglobulin genes, HLA class I and II genes and 3 specific genes with variable/noisy expression: MALAT1, JCHAIN and XIST.

To minimize technical variability, scRNAseq data from patient 5 samples, which were generated using Chromium V1 chemistry and showed significant systematic biases, were not used to generate the clustering model in steps 1–6. Patient 5 data were well projected accurately onto the model generated with the V2 chemistry and showed high similarity to data of other patients.

Cells associated with two clusters (#23 and 44) contained marker genes accounting for distinct cellular lineages and were excluded as they likely corresponded to doublets. One cluster of PC (#9) had a similar expression profile to other plasma cell clusters but was excluded as very few cells were projecting in it.

Batch corrected gene expression profiles are obtained by subtracting the expected number of UMIs associated with background noise (“noise UMIs”) from the observed number of UMIs per cluster. Gene expression profiles per cellular subtypes previously defined by grouping clusters with high transcriptional similarities (see below) were estimated using the same approach (**Table S7**).

### Gene modules

We downsampled the cells to 2000 UMIs/cells and selected variable genes similarly to the seeding step of the clustering. In order to focus on biologically relevant gene-to-gene correlation, we calculated a Pearson correlation matrix between genes for each sample and averaged the matrices following z-transformation. The averaged z matrix was then transformed back to correlation coefficients. We grouped the genes into gene “modules” by hierarchical clustering.

### Differential expression analysis

We tested for differential expression between two sets of cells by estimating the gene expression per set (similarly to the estimation of the model multinomial parameters) and calculated the observed log-fold-change between the two sets for each gene. We then randomly shuffled the cells of the two sets for 10^5^ permutations while maintaining the sizes of the sets and calculating the log-fold-change between the permuted sets for each permutation. The empirical p-value was then defined as based on the rank of the absolute value of observed the log-fold-change of each gene within its empirical fold-change distribution.

When comparing between groups of samples, we only tested genes with at least 100 cells with non-zero UMI counts and capped the number of cells per sample to equally represent the different samples within each group. Specifically, we used 50 cells/sample for the Tregs differential expression analysis between the inflamed samples of the Inf.MNP-high and Inf.MNP-low groups of patients. For the analysis the compared the total gene expression between the two patient groups we used a maximum of 1000 cells/samples.

### Characterization of lymphoid and stromal scRNAseq clusters

#### T cells characterization

Sixteen scRNAseq clusters (#17, 43, 19, 29, 22, 38, 21, 26, 4, 47, 34, 36, 30, 46, 48, 42) of T cells were characterized by shared co-expression of T-cell receptor (TCR) coreceptor encoding genes from the CD3 complex like *CD3D*, TCR costimulatory molecule *CD2* and maturation marker *CD7* (**Figures 1B and S3B**). Two clusters of innate lymphoid cells (ILCs) also expressed high levels of CD7, together with CD2, but no expression of CD3 genes or other lineage gene such as *MS4A1* (CD20), *CD19, SDC1, TNFRSF17, LYZ*, HLA-DRs, *IL3RA, TPSAB1*, as previously reported (Lim et al., 2017). Gene modules within T cell clusters were defined based on genes with strong gene-to-gene correlation of expression, as described above **(Figure S3B; Table S8**). Based on the module analysis we defined gene expression programs related to **cell cycling** (STMN1, CLSPN, CKS1B, NUF2, ASPM, UBE2T, CENPF, RRM2, DTYMK, SGOL1, CENPE, MAD2L1, CCNA2, CENPU, CCNB1, DHFR, HMMR, MXD3, GMNN, CENPW, MCM7, EZH2, SMC2, CCDC34, CDCA5, H2AFX, ZWINT, CDK1, HELLS, MKI67, TMEM106C, CDKN3, NUSAP1, CCNB2, KIAA0101, PRC1, CENPN, CDT1, AURKB, TOP2A, TK1, BIRC5, TYMS, CDC25B, PCNA, TPX2, UBE2C, ASF1B, GTSE1), **immunoregulation** (TNFRSF18, TNFRSF4, TNFRSF9, TNFRSF1B, AC133644.2, CTLA4, TIGIT, ICA1, RHBDD2, MAGEH1, IL2RA, TBC1D4, BATF, IKZF2, FOXP3), **naïve/central memory (CM)** (SELL, LEF1, SOX4, SC5D, CCR7, TOB1, NOSIP), **CD8/cytotoxic** (CD8A, CD8B, DHRS3, GZMA, GFOD1, IFITM3, PRF1, KLRD1, GZMB, CCL5, NKG7, FGR, CD160, FCER1G, XCL2, XCL1, GNLY, EOMES, CMC1, DTHD1, AOAH, CLIC3, CTSW, KLRF1, KLRC2, KLRC1, PTGDR, MCTP2, CCL3, CCL4, CCL3L3, CCL4L2, MATK, MAPK1, IL2RB) and **resident memory T cell** (JUN, KLF6, FOSB, PTGER2, FOS, SYTL3, SPRY1, ANKRD28, GPR171, PDE4D, JAML, IL7R, GLIPR1, CD69, NFKBIA, PPP1R15A, NFKBIZ, TNFAIP3, PTGER4, ANXA1, ID2, ATF3, MGAT4A, AC092580.4, KLRB1, RORA, IL18R1, STAT4, IFNGR1, PFKFB3, GPR65) states.

The total UMI counts of each of the five transcriptional programs were calculated for each cell and divided by the total number of UMIs/cell. The five scores were log transformed following division by their 75% percentile. These scores demonstrated the different distributions of the expression programs across T cell states (**Figure 3A**).

T cell cluster #17 expressed a strong cell cycle score and genes encoding for membrane proteins associated with human T cells activation such as *HLA-DRA, CD38* and the high affinity IL-2 receptor chain *IL2RA* (CD25), and thus was annotated as highly activated T cells. Cells in this cluster included both CD4 and CD8 T cells. Concordantly, these cells expressed genes associated with CD4^+^ T helper cells functions i.e. *CD40LG* or CD8^+^ T cells cytotoxicity (*GZMA, GZMB, PRF1*), indicating that T cells clustered according to their highly activated state over the CD4/CD8 lineage specification. Accordingly, both CD4^+^ and CD8^+^ HLA-DR+ activated T cells were identified by CyTOF analysis (**Figures S3C-D**).

T cell cluster #21 expressed high levels of the cytotoxic program including granzymes and perforin encoding genes, as well as low levels of the resident memory, naive/CM and cell cycling programs, and low expression of the IL-7 receptor gene *IL7R* (CD127), and thus was referred to as effector memory cytotoxic T lymphocytes (CTLs) (Mahnke et al., 2013), while T cell cluster (#38) expressed lower level of the CD8/cytotoxic program together with strong expression of genes of the γδ TCR-like *TRDC*, and was annotated as γδ T cells (**Figures 3A and S3B**).

T cell cluster #43 expressed high levels of Treg-associated genes including *FOXP3, IL2RA* (CD25), *CTLA4*, together with genes which high expression was consistently associated with Tregs by recent scRNAseq studies, including *TNFRSF18, TNFRSF4*, and *LAYN* (Savas et al., 2018; Zemmour et al., 2018; Zheng et al., 2017a) and *BATF*, recently identified as a critical TF for tissue Tregs (Hayatsu et al., 2017) (**Figures 3A and S3B**), and was thus referred to as Tregs. Interestingly, Tregs strongly expressed the co-inhibitory receptor *TIGIT*, which was shown to inhibit pathogenic Th1 and Th17 responses during murine intestinal inflammation (Joller et al., 2014), as well as the co-stimulatory receptor CD27, recently suggested as an important repressor of Th17 function acquisition by Tregs during inflammation (Rosenblum MD and coll., unpublished).

Cells in cluster #22 shared several genes with Tregs including *TIGIT, BATF, ICA1* and *TNFRSF4* (OX-40), but lacked Tregs-defining transcripts *FOXP3, IL2RA*, TNFRSF18 and *IKZF2* (HELIOS) while they expressed high levels of *CD40LG* and the chemokine *CXCL13*, and thus were referred to as T follicular helper-like (TFH-like) cells (Rao et al., 2017) (**Figure S3B**).

Two cell clusters (#19, 29) expressed high levels of the naïve/CM signature, including the chemokine receptor *CCR7* and the selectin CD62L (*SELL*), known to promote the recruitment of naïve and central memory (CM) T cells to the T cell areas of secondary lymphoid organs through the high endothelial venule (Förster et al., 2008) (**Figures 3A and S3B**). In agreement with the naïve and/or CM state of T cells in these clusters, high expression of IL-7 receptor *IL7R* (CD127) was also detected (Mahnke et al., 2013).

The first of these two clusters (#19) exhibited higher levels of the TF *LEF1*, preferentially expressed by naïve T cells, before antigen exposure (Willinger et al., 2006) and low-to-no expression of genes associated with CD4 and CD8 effector T cell functions or tissue-directing chemokine receptors. As for activated T cells, CD4^+^ and CD8^+^ T cells were mixed in this cluster, supporting dominance of the naïve state at driving the clustering, an observation already reported for blood naïve T cells (Zheng et al., 2017b). In contrast, T cells in cluster #29 expressed additional chemokine receptors like *CCR6*, together with the lectin *KLRB1* (CD161), inflammatory cytokines and chemokines (*IFNG, TNF, CCL20, CCL5*), IFNγ-induced genes (CD74, HLA-G, ISG15, IFITM1) and genes associated with inflammatory responses, thus suggestive of a population of pro-inflammatory Th1/Th17 CM T cells (Ramesh et al., 2014) (**Figures 3F, S3B-C**).

Nine clusters (#26, 4, 47, 34, 36, 30, 46, 48, 42) shared strong expression of the tissue resident memory T cells (Trm) program, which included the tissue-retention molecule *CD69*, the TFs like *ATF3, FOS, FOSB, KLF6* and the integrin *ITGA1* (CD49a), and the TF *PRDM1*, previously shown to be associated with gut Trm (Hombrink et al., 2016; Kumar et al., 2017; Mackay et al., 2016; Thome et al., 2015; Wong et al., 2016). In support of the resident memory state of these cells, the Trm clusters accounted for the large majority of the T cells in uninflamed tissues. Additional expressed genes included the prostaglandin E2 receptor (*PTGER4*)*, ANKRD28, SPRY1*, G-protein coupled receptor *GPR171*, the TF *ID2, TNFAIP3*, an NF-κb inhibitor induced by TNF protecting T cells against TNF-induced apoptosis and the *IL18* receptor subunit IL18RAP, none of which were previously described in human intestinal Trm. Five clusters (#26, 4, 47, 34, 36) corresponded to CD8 Trm. Two of these clusters were highly enriched for the cytotoxic program (#26, 4), thus reminiscent of the Trm poised for cytotoxic functions recently described in the skin (Cheuk et al., 2017). Remaining Trm clusters (#30, 46, 48, 42) expressed multiple genes associated with Th17 T cells, (*KLRB1, CCL20, IL17, IL22, MAF*, CCR6),high levels of *CD40LG* (CD154) and the cytokine IL-2, thus suggesting antigen-dependent activation (Chattopadhyay et al., 2005). The presence of naïve, CM, resident memory, highly activated and regulatory T cells in inflamed CD tissues was validated by CyTOF analysis, which also provided better granularity for CD4 and CD8 populations of naïve, CM and highly activated T cells (**Figures S3C&D**). For simplicity and based on their high transcriptomic similarities, the five clusters of CD8 Trm (#26, 4, 47, 34, 36) were pooled and referred as CD8 Trm, and the four other clusters were pooled and referred as Trm (#30, 46, 48, 42) in analyses involving T cell subtypes frequencies (**Figures 2I, 3B, 3E-I**).

### Innate lymphoid cells characterization

The two ILC clusters (#35, 28) expressed *KLRB1* (CD161), *TYROBP* (DAP12) and *FCER1G* and lacked transcript of genes specific for ILC2, like *PTGDR2, IL17RB, IL1RL1, HPGDS* or *HPGD*, and thus were referred to as Group 1 ILC1/NK and ILC3 subtypes. Cluster #35 displayed a gene signature highly compatible with ILC3, the most abundant ILC population in human healthy gut (Simoni et al., 2017), including *KIT* (CD117), *IL23R, IL1R1*, the TFs *ID2, RORC* (RORγt), *RORA* (RORα) and the cytokines *IL22* and *CSF2* (GM-CSF) (Björklund et al., 2016; Mortha et al., 2014). This cluster also expressed lower level of CD45 (*PTPRC*), a well-defined feature of both mouse and human ILC3 (Walker et al., 2013). Cluster #28 expressed cytotoxic genes (*GZMs, PRF1*), the cytokines XCL1, XCL2 and IFNγ, and CCL3/4/5 chemokines, as well as additional NK cell markers including *IL2RB* (CD122). Because some cells also expressed *IL7R*, we chose to annotate this cluster as Group 1 ILCs to encompass NK cells, CD127+ ILC1 and potential IFNγ-producing “ex-ILC3s” which origin remains unclear (Bernink et al., 2013; Simoni and Newell, 2017). The presence of ILCs was confirmed by CyTOF analysis in CD tissues but our Ab panel did not allow clear separation between the two subtypes defined by scRNAseq (data not shown).

### B cells characterization

The B cell compartment was largely dominated by plasma cells (PC) clusters (#27, 13, 20, 1, 45, 49, 50, 7), which expressed genes encoding PC-associated membrane proteins including CD138 (*SDC1*), CD38 and CD27 as well as genes associated with protein folding, trafficking and secretion (*SSR4, FKBP2, FKBP11, HSP90B1, SEC11C*), degradation of misfolded protein (*SDF2L1, DERL3*) and endoplasmic reticulum stress, including the IBD susceptibility TF gene *XBP1*. PC also expressed *LGALS1*, encoding for galectin-1, a downstream gene of the PC TF BLIMP-1 important for PC differentiation, survival and Igs secretion (Anginot et al., 2013; Tsai et al., 2008). These clusters were referred as PC expressing IgA (#27, 13, 20, 1), IgM (#45) and IgG PC (#49, 50, 7). We did not consider light chain isotype differences to annotate the clusters (**Figures S3F**). In agreement with a previous study in colonic IBD, IgG PC expressed CXCR4, the receptor for the chemokine CXCL12 (Uo et al., 2013), while IgM and IgA PC expressed higher levels of BCMA (*TNFRSF17*), the receptor for a proliferation-inducing ligand (APRIL) and B cell activating factor of TNF family (BAFF), important for PC survival in the gut (Barone et al., 2009; Castigli et al., 2004). We validated the increased levels of IgA− CXCR4_+_ PC in inflamed tissues by CyTOF analysis (**Figure S3G**). IgM and IgA-producing PC also expressed high levels of the J-CHAIN encoding gene (*IGJ*), necessary to assemble multimeric IgA and IgM and their epithelial transcytosis and secretion into the intestinal lumen through binding to the poly-Ig receptor (Spencer and Sollid, 2016). Along with PC, a small cluster of plasmablasts (#37) was identified, which together with genes involved in immunoglobulin expression and secretion described above, also expressed cell cycle genes. Aside from PC and plasmablasts, three clusters of B lymphocytes (#33, 14, 39) were identified (**Figure S3F**). B cells in these clusters shared the expression of important B cell genes including CD19, CD20, BCR co-receptors CD79a and CD79b, BANK1, MCH-II encoding genes and the TFs IRF8 and PAX5 for instance. Differential enrichment of co-expressed genes reflecting naïve (*IGHD, FCER2, CD72*) (#33) or switched memory (*CD27, IGHG1*) (#14, 39) existed between the clusters (Klein et al., 2003). As for T cells, in analyses involving PC and B cells subtypes frequencies, the clusters of IgA- and IgG-producing PC were pooled, grouping together clusters expressing different subclasses of a main class, and referred as IgA PC (#27, 13, 20, 1) and IgG PC (#49, 50, 7) respectively; the two clusters enriched in memory B cells were pooled and referred as memory B cells (#14, 39) (**Figures 2I, 3B, 3E-I**).

### Stromal and glial cells characterization

Non-hematopoietic cells identified by the scRNAseq clustering analysis included three clusters of endothelial cells (EC) (#6, 18, 12) sharing expression of the vascular endothelial growth factor receptor 2 (*KDR*) (**Figure 3C**). EC separated into one cluster of lymphatic EC (#12), uniquely defined by high levels of the lymphatic vessel endothelial hyaluronan receptor 1 (*LYVE1*) and the CCR7 ligand *CCL21*, while clusters (#6, 18) were referred to as blood EC based on co-expression of *PECAM1* (aka CD31) and Von Willebrand factor *VWF*. EC in cluster #18 expressed high levels of the atypical chemokine receptor 1 (*ACKR1*) and adhesion molecules encoding genes *SELE* (E-selectin) and *SELP* (P-selectin), suggesting they were activated (Aird, 2007), while EC in cluster #6 expressed high levels of the thrombospondin receptor *CD36*. Two clusters of mural cells (#10, 11), which are contractile cells lining the endothelium, were identified by their high expression of *MYH11, ACTA2* (αSMA) and *MCAM* (CD146). The unique expression of *CSPG4* (NG2) and *RGS5* by cells in cluster #10 identified them as pericytes (Turley et al., 2015), whereas cells in cluster #11 corresponded to smooth muscle cells, as indicated by their high expression levels of *ACTG2* (actin gamma-2 enteric smooth muscle) and *DES* (desmin).

We also identified two clusters of fibroblasts (#3, 5), characterized by shared expression of *PDGFRA, PDGFRB*, encoding for the two subunits of the platelet-derived growth factor receptors, the prostaglandin-H2 D-isomerase *PTGDS*, and several genes encoding extracellular matrix (ECM) proteins like lumican (*LUM*) and ECM remodeling proteins (e.g. matrix metallo-proteinase-2; *MMP2*). Cells in one of the two fibroblast clusters (#5) expressed a hallmark activation program including strong expression of *THY1* (CD90), *PDPN* (podoplanin), *CHTCR1* and *CHI3L1* (Turley et al., 2015), together with pro-inflammatory chemokines including neutrophils attracting *CXCL2, CXCL8, CXCL1* and *CXCL5*, as well as blood monocytes recruiting *CCL2* and *CCL7*. The last cluster of non-hematopoietic cells (#16) corresponded to *S100B* expressing neurons (Joseph et al., 2011).

### Ligand-Receptor analysis

We curated a list of cytokine/chemokine and receptor pairs and studied the association of individual ligands with the frequencies of the different cellular subtypes. Since almost all selected ligands can be secreted from expressing cells, the best estimator for the ligand concentration in the tissue given these data would be the total ligand UMI fraction across all cellular subtypes. When correlating between the expression of a specific ligand and the frequency of a particular cellular subtype, we subtracted the UMI contribution of this subtype from the total UMI counts of the ligand, since these UMI counts also depends on the size of the cell subpopulation. Specifically, we calculated the total gene expression excluding the contribution of subtype *s* of ligand *l* in sample *m* as:

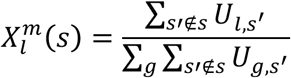

and the correlation C between ligand l and cellular subtype s as:

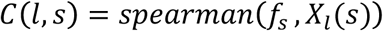

where U is the UMI matrix and *f_s_* corresponds to the normalized frequencies of subtype *s* (i.e. subtype frequency is divided by the total frequency of its cellular compartments).

### Defining the Inf.MNP module high and Inf-MNP-module low gene sets

We sought to define a scoring method that could capture the two modalities observed in our scRNA-seq ileum dataset in bulk RNA-seq or microarray data from ileal biopsy samples. We screened for genes with differential total expression between scRNA-seq inflamed samples of Inf.MNP-high or Inf.MNP-low groups and from these two lists we selected only genes that showd higher expression in cellular subtypes associated with the corresponding group. These criteria provided Inf.MNP-module associated or not-associated lists of genes differentially expressed on the bulk level in the scRNA-seq dataset.

This analysis the Inf.MNP-module-high group included patients {5,7,8,11,12}, while the Inf.MNP-module-low group included patients {10,13,14,15}. Based on the scRNA-seq results we defined two sets of cellular subtypes enriched in Inf.MNP-module-high and Inf.MNP- module-low patient groups respectively, as follows:

SubtypeSet^high^={infl.macs, mature DC, pDC, IgG-plasma cells, central memory T cells, cytotoxic T cells, Tregs, ACKR1+ endothelial cells, activated fibroblasts}

SubtypeSet^low^={resident macrophages, moDC, IgA-plasma cells, ILC3, CD8 TRM cells, TRM, cytotoxic CD8 TRM, enteric neurons, CD36+ endothelial cells, fibroblasts}

We first ran a differential expression analysis (as described above) between inflamed samples of the two groups of patients defined above, and calculated the maximum expression across all cellular subtypes in SubtypeSet^high^ and SubtypeSet^low^ for each gene, denoted m^high^ and m^low^ respectively.

We then screened for genes for which all the following conditions were true:

1. Frequency of 10^−6^ or higher in at least one of the patient groups
2. 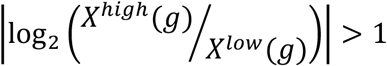 and FDR adjusted (BH) p-value <10^−3^
3. 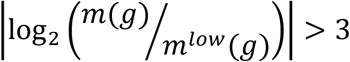

where *X*^high^ (*g*), *X*^low^(*g*) are the UMI fraction of gene g in Inf.MNP-module-high and Inf.MNP-module-low groups respectively, as provided by the differential expression analysis. Gene g was assigned to gene Set^high^ if *X*^high^(*g*) > *X*^low^(*g*) or to Set^low^ otherwise.

### External cohorts and definition of response to anti-TNF therapy

RNAseq data from the RISK cohort (n=199) were generated from ileal biopsies of children and adolescents younger than 17 years old, newly diagnosed with ileal CD and with baseline colonoscopy and confirmation of characteristic chronic active colitis/ileitis by histology prior to diagnosis and treatment, as previously described (Haberman et al., 2014).

Microarray data from the CERTIFI cohort (GSE100833) were generated at time of inclusion in the phase 2b clinical trial of Stelara® (ustekinumab) for moderate-to-severe Crohn’s disease patients with at least 3-month history of disease, and whom met the criteria for either primary or secondary non-response to anti-TNF, or had unacceptable side effects after receiving a TNF antagonist at an approved dose (Peters et al., 2017; Sandborn et al., 2012). Data from biopsies in involved ileums were analyzed (n=50).

Microarray data from the UNITI cohorts were obtained from the phase 3 clinical trial of Stelara® for moderate-to-severe Crohn’s disease (Feagan et al., 2016). Patients in the UNITI-1 cohort met the criteria for either primary or secondary non-response to anti-TNF, or had unacceptable side effects after receiving a TNF antagonist at an approved dose, while patients from the UNTI-2 cohort had failed conventional therapies but were naïve from biologics at time of inclusion. Some patients in UNITI-2 could have previously received anti-TNF but did not present with any criteria for anti-TNF non-response. Data from biopsies in involved ileums before ustekinumab introduction were analyzed (UNITI-1: n=61; UNITI-2: n=141).

### Projecting bulk RNA data onto Inf.MNP module high and Inf-MNP-module low scores

Microarray data of CERTIFI, UNITI-1, UNITI-2 were normalized using the R Bioconductor package XPS with standard parameters, applying RMA background correction, quantile normalization and farms summarization. RISK RNA-seq read counts were converted to fragments per kilobase of transcript per million (FPKM) values. Each dataset was then transformed to z-scores and projected onto the Inf.MNP-module and “No module” scores by averaging over the z-scores of genes in Set^hlgh^ and in Set^low^ respectively.

### Definition of non-response to anti-TNF

Analyses involving response or non-response to anti-TNF were conducted in patients from the RISK cohort. Inclusions were limited to patients who received anti-TNF within the first year after diagnosis. We defined strict criteria for response by considering patients achieving durable (6 months) corticosteroid-free clinical remission (pediatric Crohn’s disease index (PCDAI) <10) at months 18 and 24 post-diagnosis, as responders.

